# FIBCD1 is a Conserved Receptor for Chondroitin Sulphate Proteoglycans of the Brain Extracellular Matrix and a Candidate Gene for a Complex Neurodevelopmental Disorder

**DOI:** 10.1101/2021.09.09.459581

**Authors:** Christopher W Fell, Astrid Hagelkruys, Ana Cicvaric, Marion Horrer, Lucy Liu, Joshua Shing Shun Li, Johannes Stadlmann, Anton A Polyansky, Stefan Mereiter, Miguel Angel Tejada, Tomislav Kokotović, Angelica Scaramuzza, Kimberly A Twyman, Michelle M Morrow, Jane Juusola, Huifang Yan, Jingmin Wang, Margit Burmeister, Thomas Levin Andersen, Gerald Wirnsberger, Uffe Holmskov, Norbert Perrimon, Bojan Zagrović, Francisco J Monje, Jesper Bonnet Moeller, Josef M Penninger, Vanja Nagy

**Author notes:** Corresponding authors: Vanja Nagy, c/o CeMM, Lazarettgasse 14, AKH, BT 25.3, 1090, Vienna, Austria. Josef M. Penninger,. Life Sciences Institute, Vancouver Campus, 2350 Health Sciences Mall, Vancouver, BC Canada V6T 1Z3. Authors contributed equally (Christopher W Fell, Astrid Hagelkruys).

## Abstract

The brain extracellular matrix (ECM) is enriched in chondroitin sulphate proteoglycans (CSPGs) with variable sulphate modifications that intimately participate in brain maturation and function. Very little is known about how the changing biophysical properties of the CSPGs are signalled to neurons. Here, we report Fibrinogen C Domain Containing 1 (FIBCD1), a known chitin-binding receptor of the innate immune system, to be highly expressed in the hippocampus and to specifically bind CSPGs containing 4-O sulphate modification (CS-4S). Cultured *Fibcd1* knockout (KO) neurons lack phenotypic and transcriptomic responses to CSPG stimulation. Further, *Fibcd1* KO mice exhibit accumulation of CS-4S, likely resulting in deficits of hippocampal-dependent learning tasks and abrogated synaptic remodelling, a phenotype rescued by enzymatic digestion of CSPGs. Likewise, neuronal specific knockdown of a *Fibcd1* orthologue in flies results in neuronal morphological changes at the neuromuscular junctions and behavioural defects. Finally, we report two undiagnosed patients with a complex neurodevelopmental disorder with deleterious variants in *FIBCD1,* strongly implicating FIBCD1 in the development of the disease. Taken together, our results demonstrate that FIBCD1 is a novel, evolutionarily conserved component of ECM sulphation recognition that is crucial for neuronal development and function.

## INTRODUCTION

The brain extracellular matrix (ECM) is a dynamic microenvironment that plays a critical role in the development and maintenance of the nervous system^1, 2^. The ECM is structurally heterogenous and is composed primarily of glycans and glycoconjugates (proteoglycans, glycoproteins and glycolipids). Glycans coordinate and are essential for many neurodevelopmental processes, including axon outgrowth and guidance, synaptogenesis, migration and synaptic plasticity^2–4^. There are many inherited human disorders caused by disruptions to the glycosylation pathways. While multiple systems are often affected, most disorders also involve the central nervous system (CNS), with accompanying symptoms including congenital malformations, epilepsy, intellectual disability and developmental delay^5, 6^. Additional connections have been made between altered glycosylation, ECM composition and autism spectrum disorder (ASD), but the molecular mechanisms remain poorly understood^7^.

Proteoglycans are composed of negatively charged glycosaminoglycans (GAGs), which consists of repeating disaccharide units, covalently joined to a core protein. A seminal GAG is chondroitin sulphate (CS), a component of chondroitin sulphate proteoglycans (CSPGs). Different spatiotemporal distributions of CSPGs with variable sulphate modifications correlate with specific and discrete developmental stages as part of the dramatic reorganisation of the ECM that accompanies and regulates brain maturation^2, 8, 9^. This includes closure of the ‘critical period’ of heightened synaptic plasticity, when CSPGs condense into lattice-like structures around neurons, known as peri-neuronal nets (PNNs), which restrict juvenile synaptic plasticity and participate in memory formation, retention and extinction^1, 10, 11^.

Few cell-surface receptors for CSPGs have been identified. Receptor Protein Tyrosine Phosphatase sigma (PTPσ) and its subfamily member Leukocyte Common Antigen-Related (LAR), as well as the Nogo receptor family members, Nogo66 receptor–1 and 3 (NgR1 and 3) were demonstrated to bind CSPGs^12–14^, however, their role in the brain is unclear. Less is known regarding recognition of specifically sulphated GAGs on CSPGs. For example, CS-4,6S (or CS-E) recognition was reported for Contactin-1 (CNTN1) and in the lungs for Receptor for Advanced Glycation End Products (RAGE)^15, 16^. RAGE is also associated with Alzheimer pathology^17^. Whether CNTN1 or RAGE brain functions are linked to CS-4,6S recognition and the role of CS-4,6S in the brain is poorly understood. To date, associations between disruptions to proteoglycan receptor function and human disease have not been reported.

Here, we identify Fibrinogen C Domain Containing 1 (FIBCD1) as a novel CSPG receptor. Previous work has determined FIBCD1 to be expressed in mucosal epithelial tissues, with highest expression in the human respiratory and gastrointestinal tracts, testes, placenta and brain. FIBCD1 is a type 2 transmembrane protein with high homology to ficolins and has been shown to act as a pattern recognition receptor of chitin, found in the cell walls of fungi^18^. FIBCD1 consists of a short N-terminal cytoplasmic tail, transmembrane domain, coiled-coil region through which FIBCD1 forms homotetramers, poly-cationic region and a C-terminal extracellular fibrinogen-related domain (FReD) which participates in ligand interactions^18^. FIBCD1 was shown to limit fungal outgrowth, regulate the gut fungal mycobiome and dampen intestinal inflammation^19, 20^. FIBCD1 has also been shown to have an endocytic function and solving of the FReD crystal structure revealed potential binding sites for GAGs such as CS^21^. Despite its high expression in the brain, no function for FIBCD1 in the nervous system has been reported. Here, we demonstrate that FIBCD1 is critical for the function of the nervous system of mice and flies. Additionally, we find *FIBCD1* loss-of-function variants in two unrelated patients contributing to a complex neurodevelopmental disorder (NDD) symptomatology that includes ASD.

## RESULTS

### *Fibcd1* is expressed in the mouse brain

As the biological function of FIBCD1 in the CNS has not been explored, we first examined its expression pattern in the mouse brain. *In situ* hybridization (ISH) using complementary DNA probe pairs against *Fibcd1* mRNA in adult mouse coronal brain section revealed strong localisation of *Fibcd1* in the pyramidal cell layer of the hippocampus, granule cells of the dentate gyrus, dispersed cells of the cortex, the medial habenula and hypothalamus (Fig. 1A). Using publicly available datasets of bulk RNA seq of sorted brain cell population, brainrnaseq.org^22^, we noted *Fibcd1* expression to be highest in neurons, and virtually absent from all other cell types sequenced (Fig. S1A). While RT-qPCR of 6 different brain regions determined that some *Fibcd1* transcripts can be observed in the olfactory bulb as well, expression is highest in the hippocampus (Fig. 1B and Fig. S1B-C). *Fibcd1* transcript levels were high in embryonic mouse brain and dropped to their lowest levels at postnatal day (P) 7 and again increased during postnatal brain development to return to their high embryonic levels at P25 (Fig. 1C and Fig. S1B, D).

**Figure 1:**
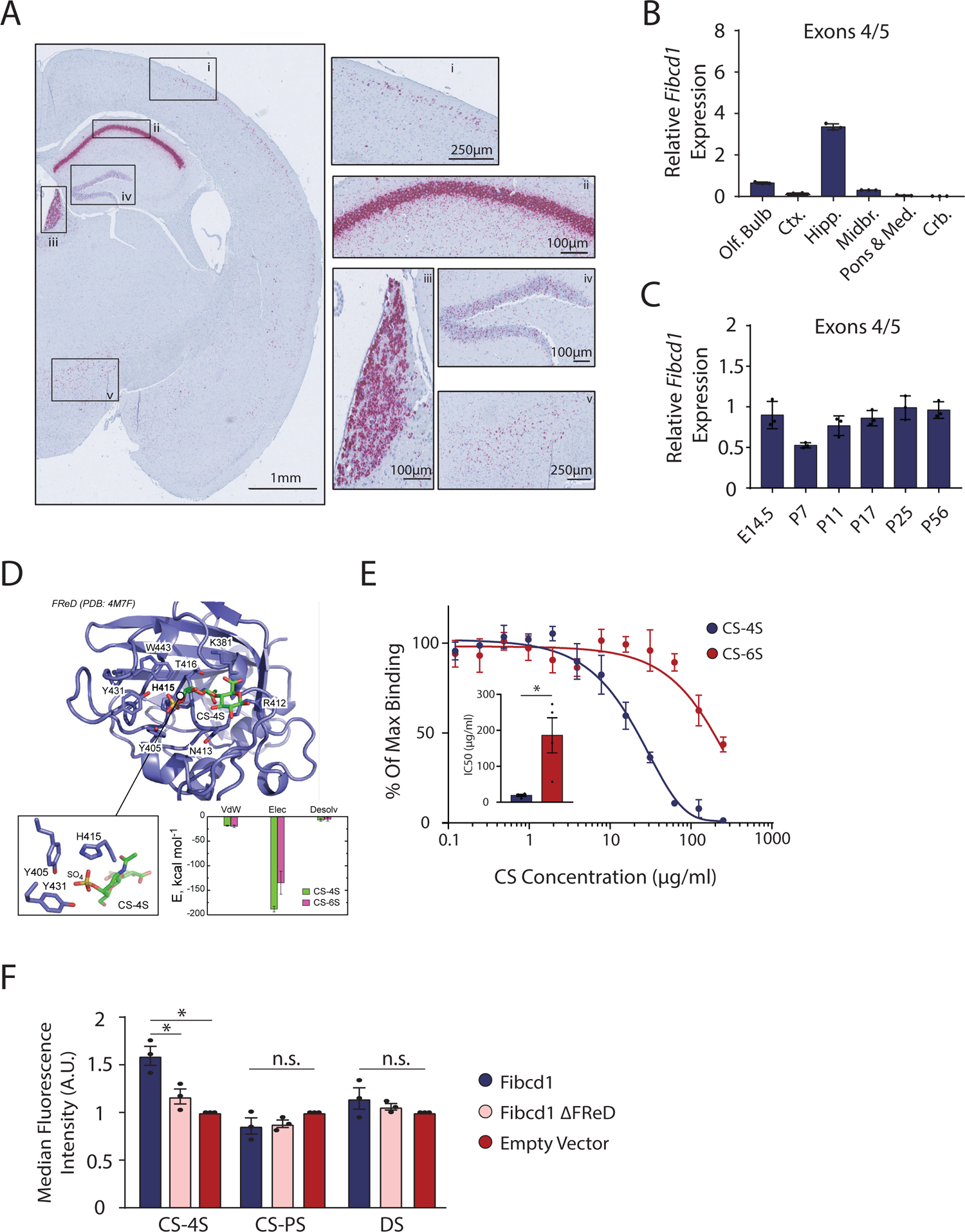
*Fibcd1* is expressed in the adult and developing mouse brain in discrete regions. (A) In-situ hybridisation with probe pairs specific to *Fibcd1* mRNA (purple) in mouse whole-brain coronal section, left hemisphere shown. Insets of high *Fibcd1* expressing regions are (i) cortex, (ii) pyramidal cell layer of hippocampus, (iii) medial habenula, (iv) granule cell layer of the dentate gyrus and (v) hypothalamus. Scale bar sizes are as indicated. Representative of 3 independent experiments. (B) Relative mRNA expression levels of *Fibcd1* (primers binding to exon 4 and 5) normalised to *Gapdh,* in the indicated adult mouse brain regions, analysed by RT-qPCR (n = 3). Olf.Bulb = olfactory bulb; Ctx. = cortex; Hipp. = hippocampus; Midbr. = midbrain; Pons & Med = pons and medulla; Crb. = cerebellum. (C) Relative mRNA expression levels of *Fibcd1* (primers binding to exon 4 and 5) normalised to *Gapdh* in the hippocampus of the indicated developmental time points, analysed by RT-qPCR (n = 3). Data is represented as mean and error bars represent SD. (D) Top binding pose for *in silico* docking of CS-4S to FIBCD1 FReD (PDB 4M7F). Inset (left) is the orientation of CS-4S within the FReD binding pocket and (right) binding free energy of CS-4S vs CS-6S. Van der Waals (vdW), electrostatic (Elec) and desolvation (Desolv) components of binding free energy charge. (E) Competitive ELISA with increasing concentrations of CS-4S (blue circles) or –6S (red circles) incubated with recombinant FIBCD1 FReD and immobilised acetylated BSA. Inset is IC50 concentrations for CS-4S and CS-6S. Data is represented as mean and error bars represent SEM. (F) Flow cytometric analysis of N2a cells expressing full-length mFIBCD1, mFIBCD1ΔFReD or empty vector control incubated with FITC-tagged chondroitin-4-sulfate (CS-4S), poly-sulphated chondroitin sulphate (CS-PS) or dermatan sulphate (DS) (N = 3). Error bars represent SEM. * = p≤0.05.

### Fibcd1 binds to chondroitin sulphate proteoglycans with a −4S modification

GAGs of proteoglycans are post-translationally modified by the addition of sulphate groups at different positions by sulfonyltransferases^23^. These include, but are not limited to, unsulphated (CS-0S), sulphated at carbon 4 (CS-4S, CS-A) and/or at carbon 6 (CS-6S, CS-C and CS-4,6S, CS-E) comprising the so-called ‘sulphation code.’ As an abundance of CS-4S and CS-6S in the brain ECM was described to be critical for maturation of the brain^8, 24, 25^, we sought to determine whether FIBCD1 binds these GAGs as previously hypothesised^21^. Top binding poses for CS-4S and CS-6S were identified using *in silico* molecular docking and an X-ray structure of the extracellular FReD (PDB 4M7F), followed by post-rescoring of docking solutions as described previously^26^. According to the scoring function, CS-4S exhibits a better fit to the FReD as compared to CS-6S (45.3 vs 43.3), with the orientations of the two ligands on the FReD surface being nearly orthogonal to each other (Fig. 1D and Fig. S2A). Importantly, the orientation of CS-4S, with its sulphate group packing tightly into a pocket formed by Y405, H415, and Y431 of FReD, leads to a more favourable electrostatic interaction and subsequently lower binding free energy (ΔΔG value of −1.3 kJ mol^-1^) as predicted by a linear model, published elsewhere^27^. To characterise binding affinities of FIBCD1 to CS-4S and CS-6S we performed competitive ELISA experiments as described previously^18^. Using a previously reported FIBCD1 ligand, acetylated BSA, and increasing concentrations of CS-4S or CS-6S, we determined a strong preference of FIBCD1 to bind CS-4S over CS-6S, with an approximately 10-fold lower IC_50_ concentration of CS-4S compared to CS-6S (Fig. 1E). To assess FIBCD1 binding specificity to CS-4S in a cellular context, we cloned V5-tagged full-length mouse WT *Fibcd1* cDNA and a truncated version without the FReD (*Fibcd1^ΔFReD^*) (Fig. S2B). We overexpressed the two FIBCD1 constructs in the mouse N2a cell line and by RT-qPCR and immunoblot analyses confirmed overexpression of FIBCD1 and V5-reactive bands at predicted molecular weights (Fig. S2C-E). We then incubated the cells with fluoresceinamine (FITC)-tagged CS-4S, polysulphated CS (CS-PS) and dermatan sulphate (DS) and acquired the cells by flow cytometry. We determined that cells expressing full length WT *Fibcd1* showed increased V5^+^/FITC^+^ fluorescence intensity in comparison to cells expressing empty vector or *Fibcd1^ΔFReD^*, but this was not the case for cells incubated with CS-PS or DS, confirming preferred binding of Fibcd1-expressing cells to CS-4S, dependent on the FReD (Fig. 1F).

We next asked whether FIBCD1 is a *bonafide* receptor for CSPGs in hippocampal neurons with high expression of *Fibcd1*. We aimed to determine cellular phenotypic and transcriptomic responses to CSPGs in primary hippocampal cultures. CSPGs are repulsive to neuronal adhesion, neurite outgrown, growth cone formation, axonal regeneration and neurogenesis in culture^12, 28^. We reasoned that if FIBCD1 is a receptor for CSPGs, then cells deficient in FIBCD1 would not exhibit the same responses as WTs as was shown for canonical CSPG receptors, such as PTPσ previously^13^. We obtained *Fibcd1* KO mice (MGI:5007144 ^29^) and validated the absence of *Fibcd1* mRNA transcript in the KO hippocampi (Fig. S3A). The mice were viable, healthy and exhibited no obvious abnormalities (Fig. S3B-D). We harvested primary hippocampal neurons from embryonic (E)18.5 *Fibcd1* WT and KO littermate animals and plated them on coverslips pre-coated with CSPGs containing a mixture of sulphated GAGs. We assessed primary neuronal morphologies after 2 and 14 days *in vitro* (DIV) by analysing cells that stain positive for the neuronal-specific marker Microtubule-associated protein 2 (MAP2). At DIV2, we found the adherence of *Fibcd1* WT neurons was partially abrogated by the presence of CSPG coating, but not in cultures plated on CSPGs after treatment with the enzyme Chondroitinase ABC (ChABC), which cleaves the GAG chains into soluble disaccharides and tetrasaccharides, leaving the core protein intact as described previously (Fig. 2A)^28, 30^. At DIV14, we found a dramatic increase in neuronal somata aggregation in cultures plated on CSPGs, also prevented by ChABC pre-digestion, again in agreement with previous literature (Fig. 2B)^28^. In contrast, neurons cultured from *Fibcd1* KO hippocampi showed slight, but not significant, DIV2 adherence impairment and slight, but also not significant, DIV14 somata aggregations, indicating that *Fibcd1* KO neurons are resistant to CSPG coating-induced phenotypes. Likely, the residual detection of CSPGs in *Fibcd1* KOs was due to the presence of other CSPG receptors, such as PTPσ^13^. Taken together, these data confirm FIBCD1 binds GAGs, particularly CS-4S, and mediates CSPG-induced cellular phenotypes.

**Figure 2:**
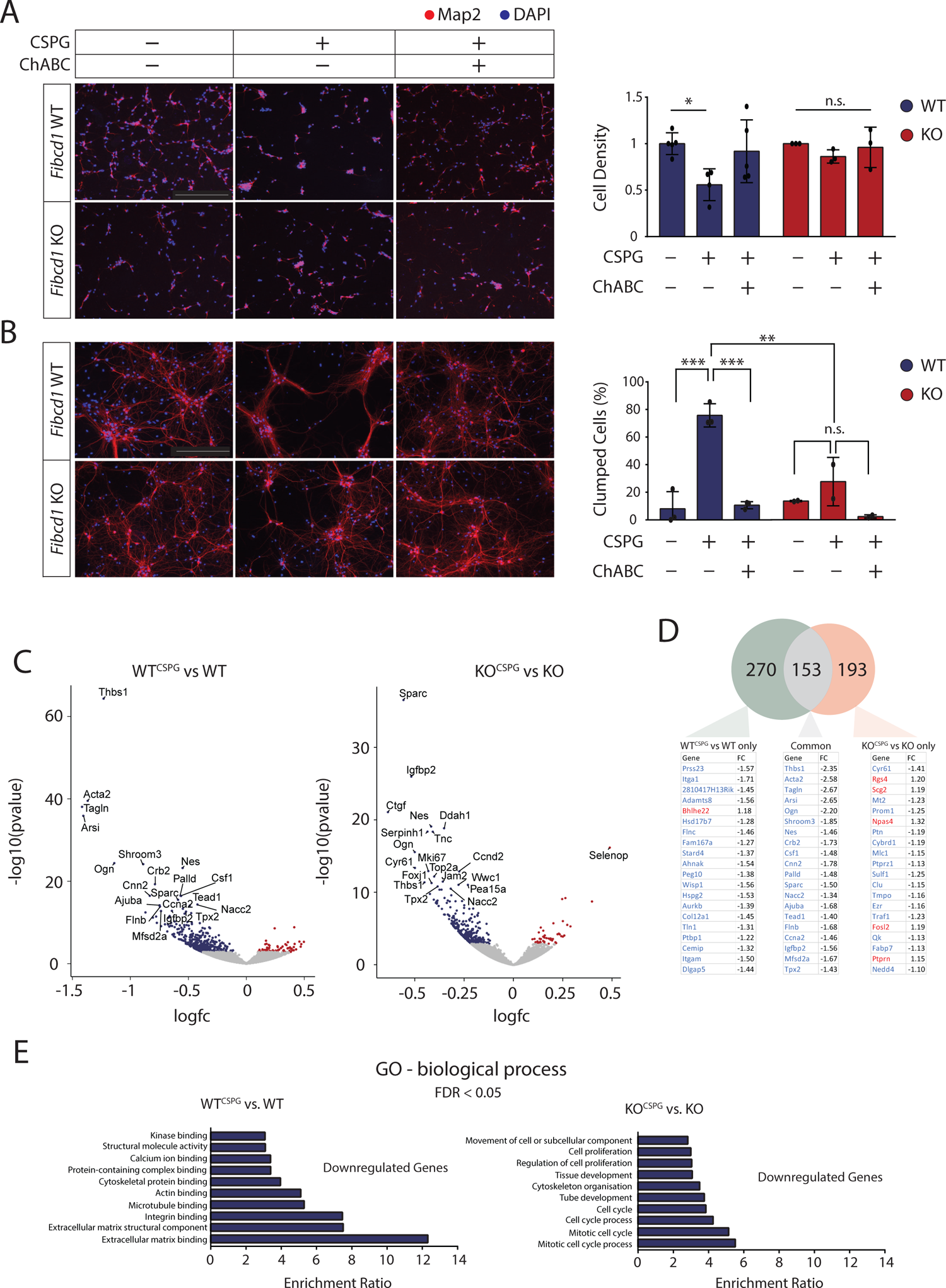
FIBCD1 mediates responses of primary hippocampal cultures to CSPGs. (A) Left, Representative images of immunofluorescent staining (MAP2, red; DAPI, blue) of primary hippocampal cultures at 2 days *in vitro* (DIV), plated on +/- CSPG coating with and without prior digestion with ChABC, as indicated. Right, quantification of DIV2 images, showing the number of protruding cells per field normalised to untreated condition. N(WT) = 3; N(KO) = 2. (B) Left, representative images of DIV14 neurons, same conditions as (A). Right, quantification of DIV14 images, representing the percentage of clumped cells per field. N(WT) = 3; N(KO) = 2. Error bars represent SD. * = p≤0.05; ** =p≤0.01; *** =p≤0.001. Scale bar = 250µm. (C) Volcano plots of differential gene expression of transcriptomes at DIV2 hippocampal cultures comparing (left) WT^CSPG^ vs WT and KO^CSPG^ vs KO (FDR < 0.05) (right). Significantly upregulated and downregulated genes are shown in red and blue, respectively. The top 20 differentially expressed genes are labelled. (D) Above, Venn diagram of significant DEGs unique to WT^CSPG^ vs WT (green, 270 genes), KO^CSPG^ vs KO (orange, 193 genes) and common between the two (grey, 153 genes). Below, lists of the 20 most significant DEGs and their fold change for each comparison, showing downregulated DEGs in blue and upregulated in red. (E) GO term enrichment analysis for significantly downregulated genes (FDR < 0.05) in (left) WT^CSPG^ vs WT and (right) KO^CSPG^ vs KO.

To investigate transcriptional responses to FIBCD1 binding of CSPGs, we isolated RNA from primary hippocampal neurons plated on coverslips coated with CSPGs (*Fibcd1* WT^CSPG^, *Fibcd1* KO^CSPG^) and without (*Fibcd1* WT, *Fibcd1* KO) at DIV3. We performed bulk RNA-sequencing with poly-A enrichment using 4 to 5 biological replicates per condition. We reasoned an early time point after plating transcriptional changes would more likely reflect cellular developmental effects of FIBCD1-CSPG binding as opposed to secondary effects such as increased cell stress, soma aggregation or dendritic fasciculation. Hierarchical clustering showed small intra-group differences and distinct separation between groups by genotype (WT or KO) and treatment (+/- CSPG) (Fig. S4A). Comparison of differentially expressed genes (DEGs, FDR<0.05) between *Fibcd1* KO and WT cells (without CSPG) revealed 462 significant DEGs with *Fibcd1* being the most downregulated DEG, as expected (Fig. S4B). We noted that a number of the top enriched DEGs in the *Fibcd1* KO vs. WT condition to be genes specifically expressed in non-neuronal cells (e.g. *Pdgfra*, *Olig2*), suggesting that DEGs may be reflecting differences between WT and KO cultures in glia numbers, which are technically challenging to control for. We therefore explored our data further comparing only between conditions within the same genotype, i.e. *Fibcd1* WT^CSPG^ vs WT and *Fibcd1* KO^CSPG^ vs KO.

Comparison between WT^CSPG^ vs WT revealed 462 significant DEGs, of which the majority (396) were downregulated. Comparison between KO^CSPG^ vs KO revealed 345 significant DEGs, of which again the majority (301) were downregulated (Fig. 2C). We cross-referenced DEGs identified in the WT and KO datasets to reveal a set of genes that are responding to CSPGs in both genotypes and those that are dependent on Fibcd1 expression (Fig. 2D). Among the top dysregulated genes common to both genotypes, independent of *Fibcd1* expression, was *Thbs1*, recently shown to be necessary and sufficient for axon regeneration after injury^31^, normally inhibited by the formation of glial CSPG scars^13^, suggesting that the effects of CSPGs may be mediated by *Thbs1* gene regulation in hippocampal neurons. Many of the remaining genes are involved in binding or remodelling of the actin cytoskeleton (*Acta2, Tagln, Shroom3, Nes, Actin, Palld, Ajuba, Flnb*) which reflect the morphological perturbations induced by plating the cells on CSPGs. Gene ontology (GO) term enrichment analysis for downregulated genes revealed terms such as “extracellular matrix binding” and “extracellular matrix structural component” in *Fibcd1* WT cells upon CSPG treatment (Fig. 2E), which suggests that Fibcd1 not only engages with the components of the ECM, but also facilitates transcriptional regulation of genes known to play a role in the ECM. Intriguingly, the third-most enriched term was “integrin binding”, reflecting a number of integrin subunits and integrin-related genes that are significantly downregulated in WT cells upon CSPG treatment (Fig. S4C). We next analysed the DEGs unique to the WT cellular response to CSPGs which are dependent on Fibcd1 expression (Fig. 2D). Among the genes dysregulated in response to CSPGs only in the WT cultures are genes coding for integrin subunits (*Itga1*, *Itgam*), integrin binding and/or modulation (*Adamts8, Tln1*)^32, 33^, genes involved in the synthesis or degradation of ECM components (*Adamts8, Hspg2, Cemip, Col12a1*)^34^ and, finally, genes involved in binding to the ECM and adhesion of cells to each other and to the ECM (*Flnc, Wisp1, Tln1*)^33, 35–38^. Together, these genes represent the transcriptional fingerprint of primary hippocampal neurons mediated by Fibcd1 binding to CSPG.

### CS-4S abundance is increased in the Fibcd1 KO hippocampus

Previous work in a non-neuronal context has shown FIBCD1 to have an endocytic function^18, 21^. Having demonstrated FIBCD1 preferential binds to CS-4S, we hypothesised that FIBCD1 may have a role in endocytosing cell-free extracellular CS-4S and/or CSPGs containing CS-4S in the brain and that a lack of FIBCD1 may cause aberrant accumulation of either or both molecules. Since CSPGs are known to condense into peri-neuronal nets (PNNs), critical for memory formation, retention and extinction^11, 39^, we first looked for any changes in PNNs in the *Fibcd1* KO brain. We found no apparent difference in the hippocampal PNNs between adult WT and KO coronal sections stained with fluorescently-tagged Wisteria floribunda agglutinin (WFA), a lectin that selectively labels CSPGs within PNNs (Fig. 3A)^40^. We also quantified amounts of sulphated GAGs (sGAG) in adult mouse hippocampi lysates, using a 1,9-dimethylmethylene blue based assay (DMMB, which reacts with sGAGs and precipitates) and found no difference in sGAG amounts between WT and KO samples (Fig. 3B). These data suggest there’s no detectable difference in general CSPG levels between *Fibcd1* WT and KO brains. To determine if particularly sulphated GAGs may be accumulating in the absence of FIBCD1 we surveyed the hippocampal glycome composition using high performance liquid chromatography (HPLC) of WT and KO mice hippocampi. HPLC analysis determined a relative increase in the ratio of CS-4S compared to other identified GAGs in the *Fibcd1* KO brains as compared to control littermates (Fig. 3C). Further, Western blot analysis using specific antibodies against differentially sulphated GAGs determined a significant increase of CS-4S abundance in the *Fibcd1*KO hippocampi as compared to controls, while CS-0S and CS-6S remained unchanged (Fig. 3D). These data indicate that while total GAG levels and overall WFA-reactive PNN-associated CSPG abundance remain unchanged, FIBCD1 deficiency leads to a specific accumulation of GAGs containing CS-4S in the hippocampus, possibly due to an inability of cells to bind and endocytose CS-4S GAGs.

**Figure 3:**
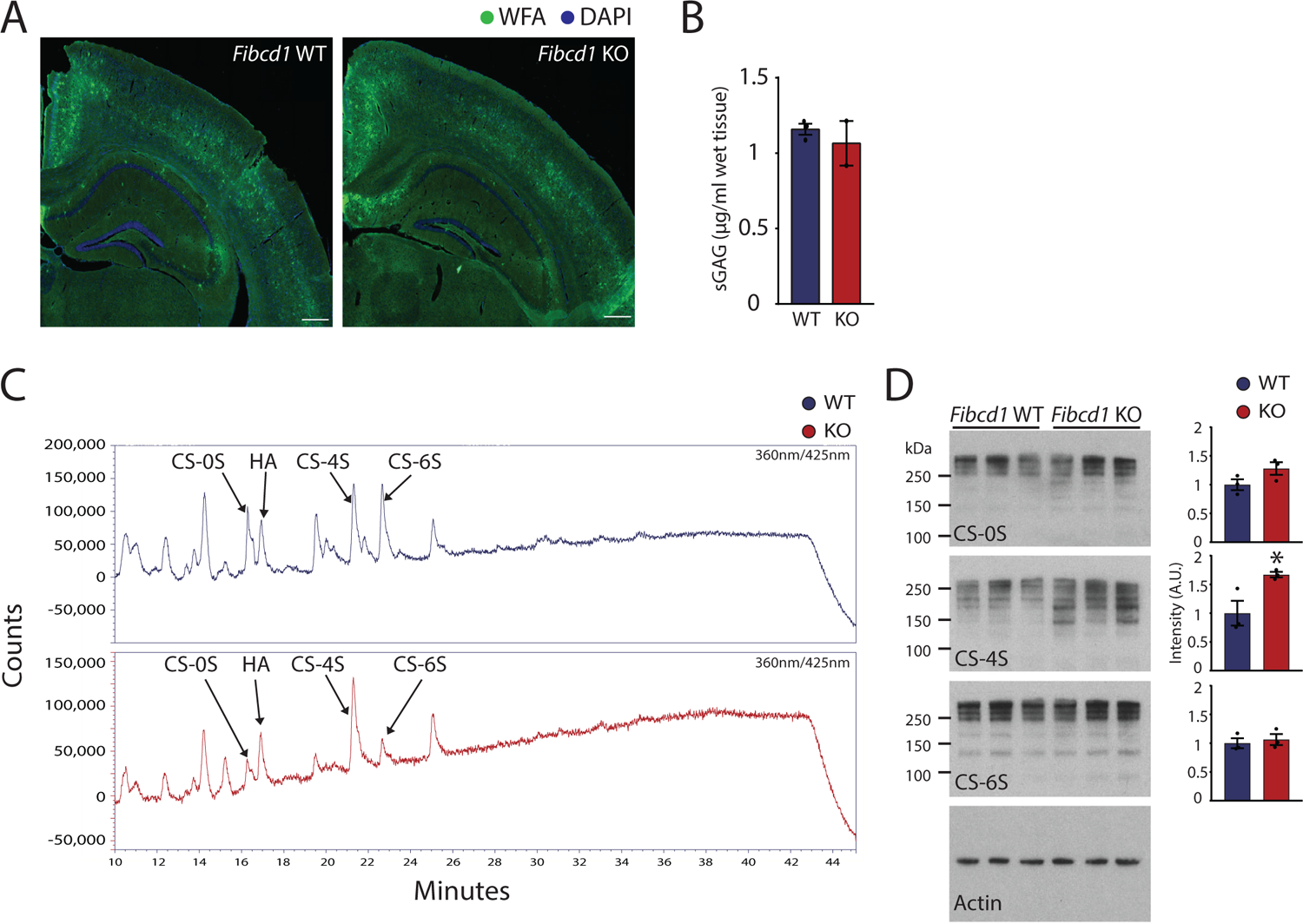
*Fibcd1* KO hippocampi have accumulated CS-4S compared to WT littermates. (A) Representative images of WFA-staining of PNNs in a coronal section of *Fibcd1* WT (left) and KO (right) mouse brains, left hemisphere (WFA, green; DAPI, blue) Scale bar = 400µm. (B) Quantification of sulphated GAG amount (per mg of wet tissue) in dissected 11-13 week old *Fibcd1* WT and KO hippocampi. Error bars represent SD. (C) HPLC traces (representative of 3 independent experiments) of variously sulphated GAGs (as labelled) in adult *Fibcd1* WT (top, blue) and KO (bottom, red) CA1 pyramidal cell layer hippocampi. Unsulphated CS = CS-0S; hyaluronic acid = HA; carbon 4 sulphated CS = CS-4S; carbon 6 sulphated CS = CS-6S. (D) Immunoblot analysis (left) and quantification of signal intensity (right) of littermate WT (blue) versus *Fibcd1* KO (red) adult hippocampi with antibodies against CS-0S, CS-4S, CS-6S and actin as a loading control. Each lane represents an independent animal (n = 3). Protein marker sizes are indicated. Error bars represent SEM. * *p* ≤ 0.05.

### Fibcd1 deficiency leads to defects in hippocampal-dependent learning and long-term potentiation in mice

To determine the behavioural consequences of CS-4S accumulation in the *Fibcd1* mouse model, we subjected *Fibcd1* WT and KO adult mice to a series of hippocampal-dependent learning tasks. Additionally, 15T MRI volumetric analysis revealed no significant differences in total brain volume, or 11 other isolated brain regions as compared to control littermates determining no overt structural abnormalities in the brains of mice with a FIBCD1 deficiency (Fig. S3D). Behaviourally, we first established that there is no difference in baseline anxiety levels between *Fibcd1* WT and KO mice as assessed by the Elevated Plus Maze (EPM) (Fig. S5A). We also detected no difference in the acquisition and retention of spatial learning between *Fibcd1* WT and KO mice, as assessed by the Morris Water Maze (MWM) (Fig. S5B-C). Nociceptive responses to noxious chemicals or heat stimulation revealed no deficiencies in sensory nervous system processing of acute pain (Fig. S5D). However, we found that *Fibcd1* KO mice were significantly impaired in spatial working memory as assessed by spontaneous alternation in the Y-Maze (Fig. 4A). Further, we found that *Fibcd1* KO animals were significantly impaired in fear-associated learning, as assessed by inhibitory avoidance (IA) task, compared to WT and heterozygous animals (Fig. 4B). Taken together, and in line with high expression of *Fibcd1* in the hippocampus, these data suggest that FIBCD1 is essential for proper mouse hippocampal development and/or function.

**Figure 4:**
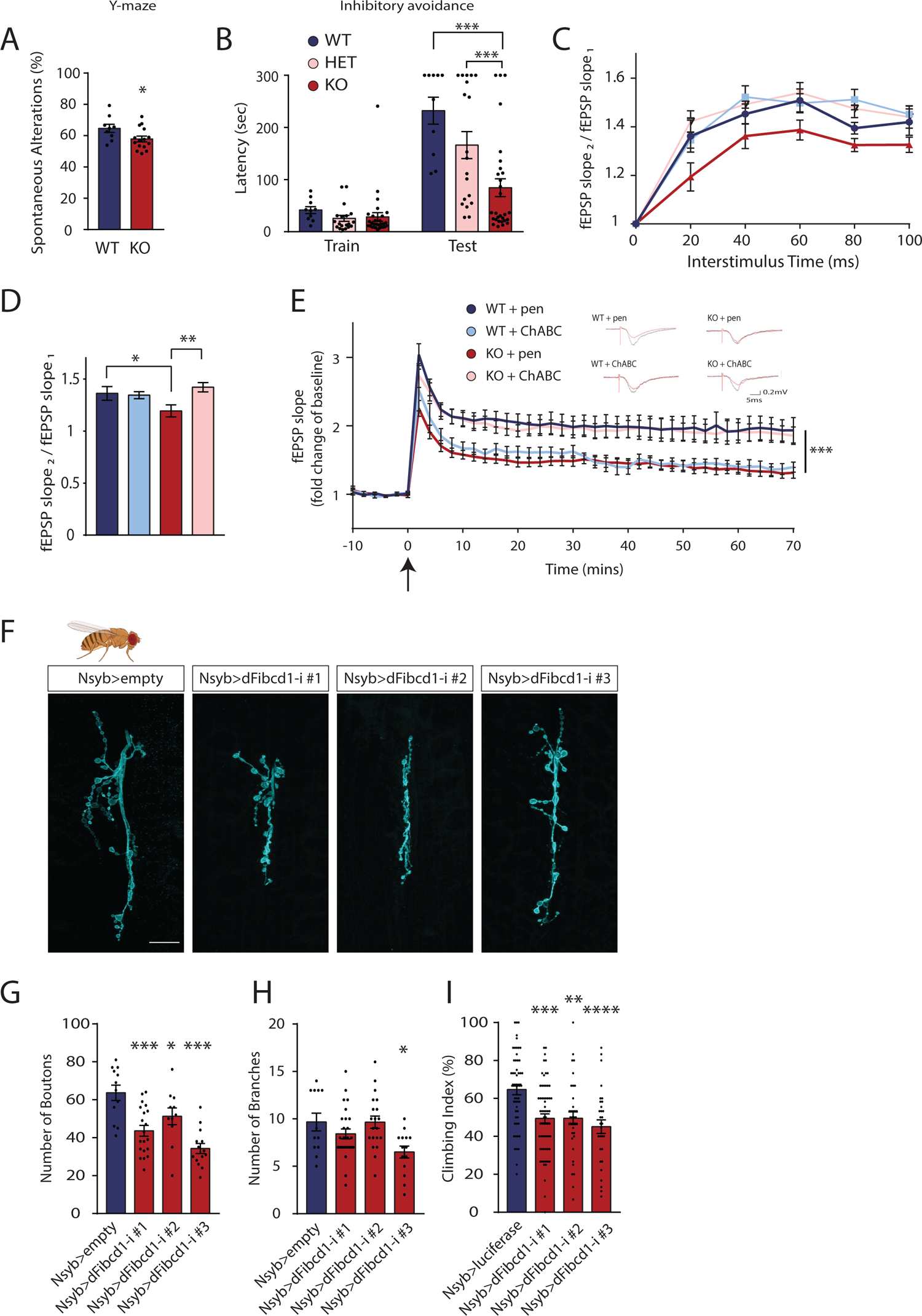
Neurological deficits in FIBCD1 deficient mice and flies. (A) Percentage of spontaneous alterations in the Y-maze. N(WT) = 9; N(KO) = 15. (B) Latency to enter the dark (foot shock) chamber during the inhibitory avoidance task at training and testing (24 hours post-training) periods. *Fibcd1* WT, heterozygous (HET) and KO mice shown. N(WT) = 8; N(HET) = 19; N(KO) = 15. Error bar represent SEM. (C-D) Paired-pulse facilitation in CA3-CA1 Schaffer collateral pathway of acute hippocampal slices from *Fibcd1* WT and KO mice. Pre-treatment with enzymes penicillinase (+pen) or Chondroitinase ABC (+ChABC) is indicated. N (WT+pen) = 17; N (KO+pen) = 20; N (WT+ChABC) = 19; N (KO+ChABC) = 25. (E) Long-term potentiation in CA3-CA1 Schaffer collateral pathways of acute hippocampal slices. Theta burst stimulation (TBS) is at time 0 indicated by the arrow. N (WT+pen) = 9; N (KO+pen) = 15; N (WT+ChABC) = 6; N (KO+ChABC) = 12. Inset are representative traces. (F) Immunofluorescent images (representative of 3 independent experiments) of control and neuronal (Nsyb) *CG10359* (*dFibcd1*) RNAi-mediated knockdown *D. melanogaster,* 3^rd^ instar larvae NMJ (NMJ6/7) stained with anti-horseradish peroxidase antibody. Empty control and RNAi-mediated knockdown of CG10359 (dFibcd1-i) lines 1, 2 and 3 shown. Scale bar = 20µm. (G) Quantification of (F), control and CG10359 knockdown lines NMJ neuron bouton number. (H) Quantification of (F), control and CG10359 knockdown lines NMJ neuron axon branch points. (E) Negative geotaxis assay of adult Drosophila control and RNAi lines #1, #2 and #3 compared to control lines expressing RNAi targeting luciferase. Climbing index represent the percentage of flies that crossed the 5 cm vial mark within 5 seconds after gentle tapping to the bottom of the vial. n is the number of tested vials. N (luciferase) = 53; N (line #1) = 63; N (line #2) = 36; N (line #3) = 31. For flies per vial, see Figure S6C). * = p≤0.05; ** =p≤0.01; *** =p≤0.001. Error bars represent SEM.

### Fibcd1 deficiency induced synaptic dysfunction are rescued by CSPG digestion

To validate our behavioural findings and ascertain the possible effects of CS-4S accumulation in synaptic function we next performed field recordings of acute hippocampal slices from adult *Fibcd1* WT and KO mice pre-incubated with ChABC or penicillinase (Pen, a treatment control that has no endogenous substrate in the brain). We hypothesised that deficiencies caused by the accumulation of GAGs containing CS-4S in the *Fibcd1* KO hippocampus could be rescued by digestion with ChABC, as is the case with other neuropathies associated with increased CSPG levels (e.g. Alzheimer’s disease^41, 42^).

Therefore, we examined the electrical properties of the CA3 Schaffer-collateral to CA1 circuit, a key pathway implicated in the formation and maintenance of spatial memories^43^. We first analysed basal properties of synaptic transmission using standard input/output protocols and found no significant differences between all conditions (Fig. S5E-F), indicating the ChABC treatment does not alter the properties of basal synaptic transmission in agreement with previous literature^44^ and indeed, no differences between pen-treated *Fibcd1* WT and KO hippocampi. We next examined paired-pulse-induced facilitation, a form of short-term pre-synaptic plasticity directly related to neurotransmitter release^45^. We observed no differences between Pen or ChABC treated WT slices, in agreement with previous literature^44^. However, slices obtained from KO mice and treated with Pen showed reduced paired-pulse-facilitation compared to Pen-treated WT slices (Fig. 4C-D). Remarkably, this reduction was restored to WT levels in the ChABC-treated KO slices (Fig. 4C-D). Finally, we examined the effects of theta-burst stimulation (TBS) induced synaptic long-term potentiation (LTP) such as the kind recorded during learning events in mice. Consistent with previous literature^44, 46^, ChABC treatment reduced, but did not abolish, potentiation in WT slices, starting at the first recorded pulse (Fig. 4E, blue traces). We found slices from KO mice pre-treated with Pen to exhibit reduced potentiation similar to WT ChABC-treated slices (Fig. 4E, dark blue vs dark red traces), but, remarkably, this deficit was again rescued in KO slices pre-treated with ChABC (Fig. 4E, pink trace and Fig. S5G-H).

### Fibcd1 neuronal function is evolutionarily conserved in Drosophila melanogaster

We next investigated if the role of Fibcd1 in the nervous system is conserved in other species. *CG10359* is a putative Fibcd1 orthologue in *Drosophila melanogaster* (*D. melanogaster*), with a high degree of FReD amino acid sequence homology (Fig. S6A) between human and mouse Fibcd1 and *D. melanogaster CG10359* (hereafter referred to as *dFibcd1*). As of yet, *dFibcd1* has no assigned function. To assess the function of *dFibcd1*, we performed RNA interference (RNAi)-mediated *dFibcd1* knockdown. We crossed three independent RNAi constructs targeting *dFibcd1* (downstream of *UAS* promoter sequence, hereafter as lines #1, #2 and #3) with lines expressing *GAL4* under the control of either the tubulin (tub) promoter for whole body RNAi expression or the *neuronal Synaptobrevin* promoter (*Nsyb*) for neuronal expression of RNAi. Full body knockdown of *dFibcd1* was lethal or semi-lethal in 2 of 3 lines (Fig. S6B), so we proceeded only with flies with neuronal knockdown of *dFibcd1*.

Neuronal knockdown of *dFibcd1* resulted in abnormal neuronal development, visualized in neuronal morphology at the larva neuromuscular junction (NMJ) (Fig. 4F). All neuronal RNAi lines exhibited reduced number of pre-synaptic boutons (Fig. 4G) and line 3 further exhibited reduced degree of neuronal branching, suggesting a reduction in neuronal function (Fig. 4H). To assess if these developmental defects also manifested as neurological phenotypes in adults, we assessed the climbing behaviour of these flies by negative geotaxis assay. We found that neuronal knockdown of *dFibcd1* resulted in reduced climbing ability when compared to control flies expressing control RNAi targeting luciferase (Fig. 4I and Fig. S6C). Taken together, our results suggest a critical role for Fibcd1 in *D. melanogaster* survival and neuronal development and function.

### Characterisation of human FIBCD1 expression and function

We next asked whether the expression and function of *Fibcd1* is conserved in humans. We profiled *FIBCD1* expression using a cDNA array from 48 different human tissues and determined that the brain is the third highest *FIBCD1* expressing tissue in humans (Fig. 5A). A further *FIBCD1* gene expression analysis of cDNA array from 24 different human brain regions determined that *FIBCD1* expression is highest in the hippocampus, followed by the hypothalamus and cortex, very similar to the *Fibcd1* expression profile in the mouse brain (Fig. 5A, inset). Considering FIBCD1’s function in mouse brain and the similarity of expression of *Fibcd1* transcripts in human and mouse CNS, we reasoned that patients with deleterious variants in *FIBCD1* would suffer from neurological dysfunctions. To our knowledge, there are no genetic disorders associated with variants in FIBCD1 to date. The Genome Aggregation Database (gnomAD^47^) that documents sequenced human genetic variants, reports 266 variants annotated as missense, stop-gained, frameshift or splice disrupting within the *FIBCD1* gene (Fig. S7A). As gnomAD does not report on zygosity or health status of any sequenced individual in the database, we searched for undiagnosed rare disease patients with high Combined Annotation Dependent Depletion (CADD) scoring variants in *FIBCD1* within our network and through GeneMatcher^48^ and were able to locate 2 unrelated patients with undiagnosed neurodevelopmental disorders. Patient 1 (P1) is a 12 year old nonverbal male from a Caucasian non-consanguineous family (Fig. 5B), with severe autism spectrum disorder (ASD), delayed verbal cognition, anxiety and attention deficit hyperactivity disorder (ADHD). He has high pain tolerance, fine motor coordination deficits and mild facial dysmorphia. Additionally, he experiences frequent allergic rhinitis and sinusitis (Table 1). There is no history of neurological disease in the family, however, several members of the maternal family have learning disabilities. As part of his clinical diagnostic evaluation, whole exome sequencing (WES) was performed at GeneDx, USA (www.genedx.com) and identifying compound heterozygous variants in *FIBCD1* Chr9:133805421 C>T; c.85 G>A; p.(G29S) and Chr9:133779621 G>A; c.1216C>T; p.(R406C), with CADD scores of 6.832 and 25.1, respectively, and a *de novo* variant in *CSMD3* Chr8: 113933925 T>C; c.1564 A>G; p.(K522E) with a CADD score of 24.7. While *CSMD3* variants have been reported in association with neurodevelopmental disorders, most published missense variants have population data in gnomAD or internal data at GeneDX, reducing the likelihood that this variant is related to the phenotype (^49^; GeneDX, Inc. personal communication). Therefore, the *FIBCD1* variants were prioritised for further analysis. Sanger sequencing determined each variant was inherited from one of the parents (Fig. 5B). There were no other variants with confirmed association to human disease identified that would match the phenotype or inheritance pattern. Patient 2 (P2) is a nonverbal 3 year old Chinese female from a non-consanguineous family with no history of genetic neurological disease (Fig. 5B), that presented with severe neurodevelopmental disorder (NDD), delayed social and cognitive abilities and delayed sitting and walking. Magnetic resonance imaging (MRI) revealed thickened cortex, decreased white/grey matter ratio, bilateral enlarged frontal gyri and ventriculomegaly (Fig. 5C). The patient also has microcephaly and dysmorphic facial features (Table 1). Additionally, recurrent pneumonia was also noted. Clinical genetic testing was performed and revealed inheritance by uniparental disomy (UPD) with mosaicism. Homozygous variants of unknown significance were found in *FIBCD1* Chr9:133779470 G>A; c.1367C>T; p.(P456L) with a CADD score of 29, *UNC13B* Chr9:35376187; c.1531T>C; p.(C511R) with a CADD score of 28.4, and *RIC1* Chr9:5765523; c.2951C>T; p.(A984V) with a CADD score of 28.6. Variants within *UNC13B* and *RIC1* were deprioritised from further functional studies due to a lack of clinical similarities with published cases^50–52^.

**Figure 5:**
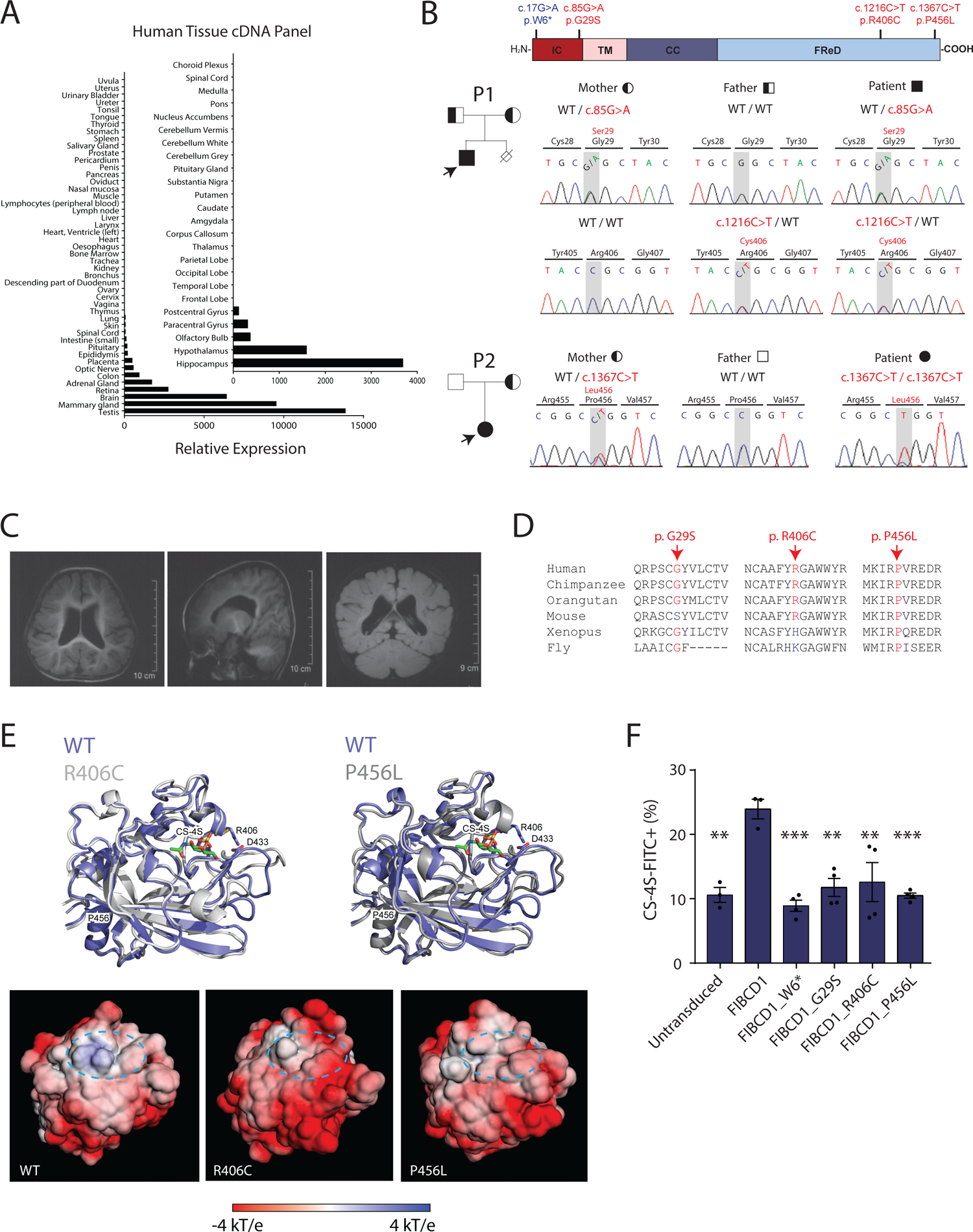
Expression of *FIBCD1 in human tissues and properties of FIBCD1* variants identified in 2 cases of undiagnosed neurodevelopmental disorders. (A) *Fibcd1* expression in various human visceral tissues and (inset) brain regions. Expression is plotted relative to the tissue with lowest detectable expression (trachea; inset, choroid plexus) (B) Top, schematic of FIBCD1 protein, with labelled intracellular domain (IC, red), transmembrane domain (TM, pink), coiled coil (CC, dark blue) and FReD in dark blue. Location of patient variants denoted in red, and of control in blue. Family pedigrees of P1 (top) and P2 (bottom) showing affected proband (filled, arrow) and carriers (half-filled, arrow). Right, representative traces of Sanger sequencing to confirm segregation within the family. P1variants are inherited in autosomal recessive manner; P2 exhibits inheritance by uniparental disomy. Above, schematic of FIBCD1 with location of patient variants and p.W6* indicated. (C) P2 MRI images (axial, coronal and sagittal plane) showing ventriculomegaly, slightly thickened cortex and bilateral enlarged gyri. (D) Amino acid sequence conservation sites of patient variants Gly29Ser, Arg406Cys and Pro456Leu in various species. (E) Top, ribbon diagrams of the superposition of the WT FReD domain with R406C (left) and P456L (right) mutants. The loops surrounding the ligand binding site (389-399 and 423-448) exhibit the largest structural rearrangement in both mutants. Bottom, comparison of the electrostatic potential mapped onto the solvent-accessible surface between WT and the two variant FReDs. (F) Flow cytometric analysis of untransduced HEK293T cells, or expressing constructs with full-length wild-type human FIBCD1, FIBCD1 with the W6* early stop variant as control (FIBCD1_W6*), or the three patient variants incubated with FITC-tagged CS-4S represented as percentage of CS-4S-FITC relative to unstained control. Statistics were calculated by 1-way ANOVA, comparing to the FIBCD1 condition. ** = < 0.01; *** = < 0.01; **** = <0.001.

**Table 1:** Summary of patient clinical findings.

|  |  | Patient 1 | Patient 2 |
| --- | --- | --- | --- |
| Background | Sex | M | F |
|  | Current Age | 12 y.o. | 3 y.o. |
|  | Ethnicity | Caucasian | Chinese |
|  | Karyogram | 46, XY | 46, XX |
|  | Consanguinity | No | No |
|  | FIBCD1 Variant | Compound Het.<br>Chr9:133805421 C>T;<br>Chr9:133779621 G>A<br>c.85G>A ; c.1216C>T<br>p.G29S ; p.R406C | Homozygous<br>Chr9:133779470 G>A<br>c.1367C>T<br>p.P456L |
|  | Family Segregation | AR | UPD with mosaicism |
|  | Other Variants | n/a | UNC13B (p.C511R) and RIC1 (p.A984V) |
| Neurology | Diagnosis | Severe ASD | Severe NDD |
|  | Psychological Evaluation | Delayed social abilities | Delayed social abilities |
|  |  | Poor eye contact |  |
|  |  | Borderline delayed cognition | Delayed cognitive abilities |
|  |  | Anxiety |  |
|  |  | ADHD combined type |  |
|  | Sitting and Walking | Normal | Delayed |
|  | Language | Nonverbal | Nonverbal |
|  | Epilepsy | - | - |
|  | Intellectual Disability | - | Too young to be evaluated |
|  | Sensory | High pain tolerance, sensitive touch | n/a |
|  | Motoric | Fine motor coordination deficits | n/a |
| Neuroradiology | MRI | n/a | Slightly thickened cortex |
|  |  |  | Decreased white matter volume |
|  |  |  | Ventriculomegaly |
|  |  |  | Bilateral enlarged frontal gyri |
|  |  |  | Microcephaly |
| Dysmorphias | Craniofacial | Triangular shaped head | Micrognathia |
|  |  | Hypertelorism |  |
|  |  | Almond-shaped eyes |  |
|  |  | Posteriorly rotated and low set ears | Low set ears |
|  |  | Epicanthal folds |  |
| Other Systems | Cardiovascular | - | Patent ductus arteriosis, resolved at 6 months |
|  | Respiratory | - | Recurrent pneumonia |
|  | Immune | Allergic rhinitis, sinusitis | - |

To determine whether the *FIBCD1* variants found in these patients affect protein folding or function we first performed all-atom molecular dynamics (MD) simulations in the microsecond range of the two *FIBCD1* variants contained within the FReD domain (p.R406C and p.P456L) and the WT as control. In the course of MD simulations, both WT and patient variant conformations stay relatively close to the initial structure, with the backbone root-mean-squared-deviations (RMSD) being the highest for R406C, intermediate for P456L and the lowest for WT (Fig. S7B), but never exceeding 2.5 Å. In order to compare WT and the two mutant structures, the dominant MD conformations were identified using structural clustering. The dominant P456L and R406C structures, deviate from the dominant WT structure by 1.6 Å and 1.5 Å backbone RMSD, respectively, while being relatively more similar to each other (1.2 Å). The largest structural rearrangements induced by the variants take place in the 389-399 and 423-448 loop regions, which surround the ligand binding site (Fig. 5E). Here, the R406C variant has a direct effect due to a disruption of the salt bridge between R406 and D433, which in the WT likely stabilises the mutual arrangement of the two loop regions. In the case of the P456L variant, the effect is rather allosteric, whereby perturbation of the conformational dynamics of the C-terminus, likely due to the removal of the sterically restricted P456, is transmitted towards the upstream 423-448 loop region. Importantly, the similar structural effect of both variants is connected to a similar perturbation of the electrostatic properties on the protein surface in the vicinity of the ligand binding site. In particular, both variants significantly increase the negative charge density of the surface patch surrounding the ligand binding site (especially R406C), in stark contrast with the WT where the corresponding surface is positively charged (Fig. 5E). We expect that this perturbation significantly weakens the binding of negatively charged phosphorylated ligands by the mutated FReDs.

To substantiate these observations, we generated HEK293T cell lines stably overexpressing FLAG-tagged human WT FIBCD1 cDNA and each of the variants G29S, R406C and P456L, as well as the W6* variant located in gnomAD database (Fig. S7A) which generates a premature STOP codon at the 6^th^ amino acid of FIBCD1, and results in a non-functional protein that serves as a negative control. We confirmed expression in each overexpressing cell line, barring W6*, by immunofluorescence (Fig. S7C). We next asked if the variants identified in our patients functionally perturbed FIBCD1:CS-4S binding, as indicated by the molecular dynamics study. We incubated HEK293T cells overexpressing WT, W6*, G29S, R406C and P456L FIBCD1 cDNAs with FITC-tagged CS-4S and acquired by flow cytometry. As with mouse WT Fibcd1 we determined that cells expressing full length human WT FIBCD1 showed increased percentage of FITC^+^ cells relative to unstained cells (Fig. 1F) or cells expressing FIBCD1_W6*. Intriguingly, this was not the case for cells expressing any of the patient variants, which exhibited similar percentage of FITC^+^ cells as the untransduced control (Fig. 5F). Together these data suggest that while the *FIBCD1* variants identified in patients did not affect protein expression, they disrupted the binding of FIBCD1 to CS-4S. Further, our molecular docking experiments suggest that, for the R406C and P456L variants, the disturbed binding may be due to a disruption of the surface electrostatic charge of the CS-4S binding pocket of FIBCD1’s FReD. Taken together, we conclude that P1 and 2 carry variants deleterious to FIBCD1 function potentially contributing to their symptoms and reveal FIBCD1 as a novel candidate gene for NDD.

## DISCUSSION

Here we report that FIBCD1, highly expressed in the mammalian brain, is a novel receptor of CSPGs, which preferentially binds to CS-4S GAGs and mediates a transcriptional program required for cell-cell and cell-matrix interactions. Using genetically modified mice and flies, we find that FIBCD1 is critical for neuronal maturation and function, whose downregulation leads to neurodevelopmental impairments and hippocampal-dependent learning deficiencies, likely resulting from aberrant accumulation of CS-4S. Additionally, we have identified 2 unrelated patients with complex NDDs that include ASD, carrying deleterious variants within the *FIBCD1* gene, implicating FIBCD1 as critical to CNS development in humans and as a novel risk gene for NDDs.

FIBCD1 is an endocytic lectin, reported to bind chitin on cellular walls of pathogens and to regulate the innate immune response to candida infections^18, 19^. We provide computational, biochemical, cellular and *in vivo* evidence that FIBCD1 also has endogenous ligands in the brain and intimately participates in the regulation of ECM composition through endocytosis or receptor-mediated signalling, or both. Indeed, transcriptomic changes upon CSPG stimulation of *FIBCD1* WT and KO primary hippocampal cultures reveal a novel ligand-dependent signalling function for FIBCD1, primarily encompassing genes involved in ECM binding and structure. Interestingly, a number of DEGs were integrin subunits or integrin-related genes, molecules well known for interacting with the ECM and signalling during neuronal development and synaptic activity^1,4^. Considering that closely related proteins containing FReDs have been shown to directly interact with integrins^53^, it is tempting to speculate a physical Fibcd1:integrin interaction, the mechanism of which remains to be uncovered. We cross-referenced the DEGs present in both the WT^CSPG^ vs WT and KO^CSPG^ vs KO datasets to identify the genes specifically regulated by Fibcd1 binding to CSPGs. We identified a number of genes coding for integrin subunits or integrin binding and/or modulation as well as genes involved in the synthesis or degradation of ECM components and, finally, genes involved in binding to the ECM and adhesion of cells to the ECM. While the functions of many of these genes have been elucidated in a non-neuronal context, it’s likely that their function is largely conserved in neurons and, therefore, these genes make up the transcriptional fingerprint regulated by FIBCD1’s interaction with CSPGs in primary hippocampal neurons.

The accumulation of CS-4S in the *Fibcd1* KO mouse hippocampi, and the ChABC-mediated rescue of LTP, strongly support the primary role of FIBCD1 as an endocytic receptor. However, it is unclear whether FIBCD1 binds entire CSPGs or its specifically sulphated GAG components and how this binding activates transcriptomic changes noted in neuronal cultures *in vivo*. The complete rescue of LTP deficits by ChABC digestion of CSPGs suggests that the hippocampus develops normally in the absence of FIBCD1 and it is the pathological accumulation of GAGs containing CS-4S over time that is responsible for the observed behavioural and synaptic phenotypes. This is in contrast to the dependence of *dFibcd1* expression in *D. melanogaster* survival and neuronal development as exhibited by the lethality of the full body *dFibcd1* knock-down flies, suggesting that *Fibcd1* may have developed a more specialised role in mammals. Nevertheless, it will be of great interest to delineate the role of CS-4S during activity-driven synaptic activity, in what context it is released from the ECM and how it contributes to hippocampal function.

It is widely recognised that brain ECM composition is critical for CNS development and function and there is growing evidence that deleterious variants in genes encoding glycoproteins, enzymes involved in glycosylation, or those required for their clearance and degradation contribute to neurodevelopmental dysfunctions, such as ASD^7^. Here, we presented evidence for an additional risk gene, *FIBCD1,* in two patients that exhibit symptoms of severe neurodevelopmental dysfunctions, including delayed social, cognitive and verbal abilities, ASD, ADHD, facial dysmorphias and structural brain anomalies. P2 was too young at last examination to be fully evaluated for ASD or ID, however, while exhibiting similar symptoms as P1, was more affected. Intriguingly, signs of immune system dysfunction such as recurring allergic rhinitis, sinusitis and pneumonia in both patients is in line with the literature describing FIBCD1 in immune responses^54^. In addition to *FIBCD1* variants, P2’s exome sequencing revealed additional variants of unknown significance (VUS) in *UNC13B* and *RIC1. UNC13B* encodes a pre-synaptic protein highly expressed in the brain, MUNC13-2, that has recently been associated with partial focal epilepsy^52^, not found in P2, and was therefore dismissed as potentially causative in this case. Variants in *RIC1* gene have recently been associated with autosomal recessive CATIFA Syndrome (cleft lip, cataract, tooth abnormality, intellectual disability, facial dysmorphism, attention-deficit hyperactivity disorder, OMIM: 618761)^50, 51^. With the exception of P2’s micrognathia, she exhibits none of the hallmark symptoms of CATIFA syndrome (cleft lip, cataracts, tooth abnormalities). However, the contribution of the RIC1 variant to the overall clinical pathology of the patient cannot be ruled out, even if unlikely.

*In silico* molecular simulation studies and cellular assays have determined the deleteriousness of the identified *FIBCD1* variants. Molecular modelling analysis has suggested that the FReD variants, R406C (P1) and P456L (P2), both lead to FIBCD1 loss-of-function by disrupting the binding pocket’s electrostatic charge and diminishing the affinity to its GAG ligand, which is consistent with our cellular assay for FIBCD1:CS-4S binding. However, it is less clear how the other P1 variant, G29S, disrupts binding of FIBCD1 to CS-4S. While we find the glycine at this residue is largely conserved among other species, the mouse orthologue contains the same substitution of glycine to serine as in P1, yet this substitution in the human *FIBCD1* orthologue was similarly disruptive of binding to GAGs as the other variants. How the function of G29 residue diverges from mouse to human, whether it is important for structural confirmation of FIBCD1, targeting, or downstream signalling remains to be elucidated. Nevertheless, we demonstrated all three FIBCD1 variants to be deleterious to protein function of FIBCD1 and in view of the data in the model organisms and cell culture, is likely to be causative of the patients’ symptoms. Full understanding of the neuropathology caused by deleterious *FIBCD1* variants awaits more clinical cases for comparison of their clinical symptoms.

Overall, FIBCD1 is a regulator of CS-4S content in the hippocampal ECM and a mediator of ECM signalling. FIBCD1 is a critical regulator of synaptic plasticity events underlying certain types of learning and memory in mice and a novel risk gene for a complex form of neurodevelopmental disorder.

## AUTHOR CONTRIBUTIONS

Conceptualisation: V.N. Formal Analysis: C.W.F., A.H., A.C., L.L., J.S.S.L., M.A.T., T.K., S.M., J.S., A.A.P., A.S., M.M.M., J.J., J.B.M., V.N. Funding Acquisition: J.P., V.N. Investigation: C.W.F., A.H., A.C., L.L., J.S.S.L, M.A.T., M.H., S.M., J.S., A.A.P., A.S., K.A.T., H.Y., J.W., T.L.A., G.W., J.B.M., V.N. Resources: N.P., B.Z., F.Q.M., J.B.M., J.M.P., V.N. Supervision: U.H., N.P., B.Z., F.Q.M., J.B.M., J.M.P., V.N. Visualisation: C.W.F., A.H., A.C., M.H., L.L., J.S.S.L., S.M., M.A.T., J.S., A.A.P., J.B.M., V.N. Writing – original draft: C.W.F., V.N., with contribution from all authors.

## ACKNOWLEDGMENTS

We thank Malene Hyggelbjerg Nielsen for excellent technical assistance with the ISH data. We are thankful to all members of the IMP-IMBA Bio-optics Core Facility for assistance in cell sorting and imaging as well as the Molecular Biology Service for their help. Moreover, we thank the Preclinical Phenotyping Facility, the Preclinical Imaging Facility and the Next Generation Sequencing Facility at the Vienna BioCenter Core Facilities GmbH (VBCF), member of Vienna BioCenter (VBC), Austria. C.W.F. is funded by a DOC fellowship of the Austrian Academy of Sciences (OeAW): 25525; L.L. is a fellow of the Damon Runyon Cancer Research Foundation (DRG-2319-18); S.M. is funded by the European Union’s Horizon 2020 research and innovation programme under the Marie Sklodowska-Curie grant agreement No. 841319; J.B.M. is funded by the Novo Nordisk foundation; J.M.P. is supported by the Austrian Federal Ministry of Education, Science and Research, the Austrian Academy of Sciences and the City of Vienna and grants from the Austrian Science Fund (FWF) Wittgenstein award (Z 271-B19), the T. von Zastrow foundation, and a Canada 150 Research Chairs Program (F18-01336); V.N. is funded by Ludwig Boltzmann Gesellschaft core funding.

## MATERIALS AND METHODS

### Patients and Whole Exome Sequencing

All procedures were performed following informed consent and approval from patients and relatives.

#### Patient 1

Using genomic DNA from the proband and parents, the exonic regions and flanking splice junctions of the genome were captured using the IDT xGen Exome Research Panel v1.0 (Integrated DNA Technologies, Coralville, IA). Massively parallel (NextGen) sequencing was done on an Illumina system with 100bp or greater paired-end reads. Reads were aligned to human genome build GRCh37/UCSC hg19, and analysed for sequence variants using a custom-developed analysis tool. Additional sequencing technology and variant interpretation protocol has been previously described^55^. The general assertion criteria for variant classification are publicly available on the GeneDx ClinVar submission page (http://www.ncbi.nlm.nih.gov/clinvar/submitters/26957/).

#### Patient 2

Procedures were in accordance with the ethical standards and approval of the Medical Ethics Committee of Peking University First Hospital. The IRB number is No. [2005]004. Patients were sequenced and analysed as described previously^56^, with sequencing performed by Joy Oriental Co. (Beijing, China).

#### Population Genetics

CADD scores and amino acid positions for FIBCD1 variants present in the population were extracted from GnomAD ^47^ and visualised using PopViz ^57^.

### Animals

#### Mus musculus

Fibcd1tm1Lex mice (MGI: 5007144)^29^ were bred on a C57BL/6J genetic background. Mouse genotypes were assessed by PCR (see Supplementary Table 1 for genotyping primer sequences). Of note, only age- and sex-matched littermates from respective crosses were used for experiments. All mice were housed at the Comparative Medicine Mousehouse Vienna BioCenter, Vienna, Austria, maintained at a 12-h light/dark cycle and provided with food and water *ad libitum*. Experiments were approved by the Bundesministerium für Wissenschaft, Forschung und Wirtschaft (BMWFW-66.009/0048-WF/V/3b/2018) and carried out according to EU-directive 2010/63/EU.

#### Drosophila melanogaster

All studies were performed in age, light, sex and temperature matched flies. All crosses were raised at 25°C on standard molasses fly food unless otherwise noted.

#### *In situ* hybridisation

The brain tissue was dissected from two 8-10 weeks old C57B6J mice and fixed in buffered 4% paraformaldehyde, dehydrated and paraffin-embedded. 3.5-µm-thick frontal sections were *in situ* hybridized with an enhanced RNAScope 2.5 high-definition procedure (310035, ACD Bioscience), as described previously^58^. The sections were hybridized with 20 probe pairs for mouse *Fibcd1* mRNA (524021, ACD Bioscience, targeting nucleotide 629 – 1567) or probe diluent as negative control. The probe pairs were branch amplified as instructed by the manufacturer, and further tyramide-enhanced using digoxigenin-conjugated tyramide (NEL748001KT, PerkinElmer) followed by alkaline phosphatase-conjugated sheep anti-digoxigenin FAB fragments (11093274910, Roche) and visualization with Liquid Permanent Red (Agilent Technologies). Finally, the sections were counterstained with Mayer’s hematoxylin.

### RNA-sequencing

RNA was isolated using Qiagen RNeasy Mini kit (74104). Library prep, sequencing and alignment were done at the Vienna BioCenter Next Generation Sequencing Facility. RNA was quantified on a Qubit Fluorometric Quantitation system (Life Technologies) and RNA integrity score calculated using a Bioanalyzer (Agilent). Poly-A enrichment was performed using NEBNext Poly(A) mRNA Magnetic Isolation Module (NEB #E7490) and library prep performed using NEBNext Ultra II Directional RNA Library Prep Kit for Illumina (NEB #E7760) and sequenced on an Illumina HiSeq 3000/4000, 50bp single-read.

### RNA-seq analysis

Demultiplexing, quality control, and alignment to the genome was done by the VBC Next Generation Sequencing Facility (Vienna, Austria). Calculation of differentially expressed genes (DEGs) was performed using Bioconductor DESeq2 package^59^. All DEGs located on the X or Y chromosomes were excluded due to uncontrollable distributions of male and female E18.5 pups from which hippocampal neurons were harvested. Sequencing data was uploaded and analysed on the Galaxy web platform (www.usegalaxy.eu)60. GO term enrichment analysis was performed using WebGestalt^61^, over-representation analysis method.

### RT-qPCR

#### Mouse

For mouse *Fibcd1* expression analysis by RT-qPCR, tissues and cells were isolated and homogenized in TRIzol reagent (Invitrogen). Total RNA was isolated according to the manufacturer’s instructions. RNA was reverse transcribed with the iScript cDNA synthesis kit (Bio-Rad). Real-time PCR analysis was performed with GoTaq qPCR master mix (Promega) on a CFX384 system (Bio-Rad). Data were normalized to values for the housekeeping gene *Gapdh.* See Supplementary Table 1 for a list of primers used in this study.

#### Human

Human cDNA panels were obtained from Origene: TissueScan, Human Brain cDNA Array (#HBRT101), TissueScan, Human Normal cDNA Array (#HMRT304). Master mixes containing primer pairs against *FIBCD1* or *B2M* (Beta-2-microglobulin, housekeeping gene) were prepared with iTaq Universal SYBR Green Supermix (Bio-Rad #1725122). The mastermixes were dispensed into each well containing cDNA and incubated on ice for 15 minutes to allow the cDNA to dissolve, then briefly vortexed. Reactions were run on Applied Biosystems StepOne Plus machine with standard settings including the melt curve. Data were normalised to values for the housekeeping gene *B2M.* See Supplementary Table 1 for a list of primers.

### *In Silico* Modelling of FIBCD1

#### Docking

*In silico* molecular docking was performed using GOLD software version 5.2.2^62^ and an X-ray structure of the extracellular FReD (PDB 4M7F)^21^. The post-rescoring of docking solutions (100 in total) was carried out as described previously^26^. The binding free energy of CS-4S and CS-6S to FReD domain was estimated using PRODIGY-LIGAND^27^ after refinement of the complexes using HADDOCK2.2 web-server^63^.

#### Patient Variant Modelling

The initial protein configuration was taken from the X-ray structure of the FReD (PDB: 4M7F; aa 239-458), where single point mutations R406C and P456L were introduced using PyMol^64^.

The WT and the two mutant structures were used for further all-atom molecular dynamics (MD) simulations in rectangular boxes (6×6×6 nm^3^) filled with ∼6000 explicit water molecules. A total of 1 µs of MD statistics was collected for each system. All MD simulations and the analysis were performed using GROMACS 5.1.4 package^65^ and Amber99SB-ILDN force field^66^. After initial energy minimization, all systems were solvated using TIP3P water model^67^. The solvated systems had neutral net charge and effective NaCl concentration of 0.1 M. These systems were again energy-minimized and subjected to an MD equilibration of 30000 steps using a 0.5 fs time step with position restraints applied to all protein atoms (restraining force constants Fx=Fy=Fz=1000 kJ mol^-1^ nm^-1^) and 250000 steps using a 1 fs time step without any restraints. Finally, production runs were carried out for all systems using a 2 fs time step. A twin-range (10/12 Å) spherical cut-off function was used to truncate van der Waals interactions. Electrostatic interactions were treated using the particle-mesh Ewald summation with a real space cutoff 12 and 1.2 Å grid with fourth-order spline interpolation. MD simulations were carried out using 3D periodic boundary conditions in the isothermal−isobaric (NPT) ensemble with an isotropic pressure of 1.013 bar and a constant temperature of 310 K. The pressure and temperature were controlled using Nose-Hoover thermostat^68^ and a Parrinello-Rahman barostat^69^ with 0.5 and 10 ps relaxation parameters, respectively, and a compressibility of 4.5 × 10^−5^ bar^−1^ for the barostat. Protein and solvent molecules were coupled to both thermostat and barostat separately. Bond lengths were constrained using LINCS^70^. Root-mean-squared deviations (RMSD) from the starting configuration were calculated using *rms* utility (GROMACS) over all backbone atoms. Conformational clustering of MD trajectories was performed using *cluster* utility (GROMACS) with the applied RMSD cut-off for backbone atoms of neighboring structures of 0.9 Å – a minimum value at which only a single dominant state was identified for WT. Electrostatic potential was calculated and mapped onto the protein solvent accessible surface using the APBS web-service (https://server.poissonboltzmann.org)71. Protein structures were visualized using PyMol^64^.

### Binding assays

Characterisation of FIBCD1 binding specificity to chondroitin sulphate 4S (CS-4S) and chondroitin sulphate 6S (CS-6S) was performed through ELISA-based inhibition experiments essentially as described previously^18^. In brief, microtiter plates (MaxiSorp, NUNC) were coated with 1 µg/ml acetylated BSA (Sigma) and blocked with TBS containing 0.05% Tween. Recombinant FIBCD1-FReD^18^ was added to each well in a constant concentration of 100 ng/ml in TBS containing 0.05% Tween and 5mM CaCl_2_ together with 2-fold dilutions of CS-4S and CS-6S (Amsbio). After incubation overnight at 4°C and washing, the wells were incubated for 2 hours at room temperature with 1 µg/ml biotinylated anti-FIBCD1 (clone 11-14-25), before washed and developed using HRP-conjugated streptavidin and TMB substrate.

### Tissue Culture

#### Primary Neurons

Timed pregnancy E18.5 pups were sacrificed by decapitation, the hippocampi dissected into 1x Hank’s Buffered Saline Solution (HBSS, Gibco #14185045) with 100 U/ml penicillin/streptomycin (Biowest #L0022-020). The tissue was minced, incubated with trypsin (0.025%) for 15 minutes at 37°C, inverting every 5 mins, and the trypsin inactivated with 10% FCS-containing media. The tissues were then washed 3 times with HBSS and triturated with heat-polished glass pipettes. 200,000 dissociated cells were plated in 1 ml of Neurobasal medium (ThermoFisher #21103049) supplemented with 10% FCS, 2mM L-glutamine (Gibco #25030149), 1x B27 (Gibco #17504001), 10mM Hepes (Gibco #15630056), 100 U/ml pen/strep in a 12-well plate and incubated at 37°C, 5% CO_2_. 50% of media was exchanged to FCS-free medium after 24 hours and then every 36 hours. Cells were plated on coverslips coated with 0-4µg/ml CSPGs (Merck #CC117) with and without prior ChABC (Sigma-Aldrich C3667) digestion as described previously^13, 28^.

### FIBCD1 overexpression

#### Mouse

Fibcd1 mouse cDNA with 3’ V5 tag was ordered as a G-block from IDT and cloned along with ΔFReD construct via restriction enzyme insertion cloning, using 5’ XhoI and 3’ EcoRI restriction enzyme sites. The constructs were cloned into a custom pMSCV-IRES-mCherry plasmid, see supplementary data for construct sequences.

Fibcd1 constructs were packaged into virus by CaCl_2_ transfecting 70-80% confluent 10cm^2^ plate of Platinum-E (PlatE) HEK 293T cells (Cell Biolabs #RV-101) with 10µg Fibcd1 construct and 2µg gag/pol plasmid. The media was exchanged next day to remove transfection mixture. N2a cells’ media was exchanged with virus-containing supernatant at 48- and 72-hours post-transfection with 8µg/ml polybrene (Sigma #H9268). N2a cells were expanded and FACS sorted for mCherry+ cells.

N2a cells stably expressing full-length V5-tagged FIBCD1, FIBCD1 lacking the fibrinogen domain (ΔFReD) or the empty vector (pMSCV-IRES-mCherry) were grown in DMEM supplemented with 10% foetal calf serum, 100 U/ml penicillin (Sigma) and 100µg/ml streptomycin sulphate (Sigma).

#### Human

*hFIBCD1* cDNA was ordered from Origene (#RC206180) and sub-cloned by Gateway cloning (Thermo) into a custom plasmid (via pDONR201) with 3’ 3xFLAG tags, mCherry and Blasticidine resistance selection markers, sequences are available in supplementary data. The PCR primers before the BP reaction were designed to remove the 3’ STOP codon present in the Origene clone to allow read-through to the 3’ 3xFLAG tags. Point variants were introduced into *hFibcd1* cDNA while cloned in the intermediate pDONR201 plasmid using the Q5 site-directed mutagenesis kit (NEB #E0554), following the manufacturer’s protocol, and the resultant clones sequenced to confirm correct base substitution before continuing to the LR reaction into the destination plasmid.

Cloned plasmids were packaged in Lenti-X 293T cells (Takara Biosciences #632180), transfected using Lipofectamine 2000 (Thermo #11668027), following the manufacturer’s protocol. Per well of a 6 well plate, 1200ng of Fibcd1 plasmid was mixed with 150µl Opti-Mem (Gibco #31985062), 700ng pGag/pol, 460ng pVSVg and 280ng pAdVAntage (Promega #E1711) and transfected for 6-8 hours. Virus-containing supernatant was collected at 48 and 72 hours post-transfection.

HEK 293T cells were transduced by spin infection using the above lentivirus and 6µg/ml polybrene and transduced clones were selected using 14µg/ml blasticidine for 48 hours. Stably transduced cells were maintained in DMEM (Thermo #11960044), 10% FCS, pen/strep and sodium pyruvate. Transduced cells were periodically re-selected with 14µg/ml blasticidine for at least 48 hours.

### Flow cytometry

#### Mouse

N2a cells stably expressing full-length V5-tagged mFIBCD1, mFIBCD1 lacking the fibrinogen domain (ΔFReD) or the empty vector (pMSCV-IRES-mCherry) were washed once with PBS and incubated for 4 hours with 100µg/ml fluoresceinamine labelled glycosaminoglycans: 4-O-sulfated chondroitin sulphate (AMS.CSR-FACS-A1, AMSBIO), poly-sulphated chondroitin sulphate (AMS.CSR-FACS-P1, AMSBIO) or dermatan sulphate (AMS.CSR-FADS-B1, AMSBIO) in DMEM. See Supplementary Table 2 for a list of flow cytometry antibodies used in this study. The staining medium was removed, cells were collected by gentle flushing with PBS and fluorescence was measured by FACS LSR Fortessa (BD). The experiment was performed in three independent replicates and analysed by FlowJo v10.6 (FlowJo LLC).

#### Human

Untransduced HEK 293T cells or those stably expressing full length C-terminally 3xFLAG tagged hFIBCD1, hFIBCD1_W6*, hFIBCD1_G29S, hFIBCD1_R406C, hFIBCD1_P456L were seeded in full media. The next day, cells were washed 1x with PBS and trypsinised for precisely 1.5 minutes, pelleted at 4°C, and resuspended in 10µg/ml 4-O-sulfated chondroitin sulphate (AMS.CSR-FACS-A1, Amsbio) diluted in freshly prepared PBS (0.8mM CaCl_2_) and incubated in a humidified incubator for 45 minutes. The cells were washed 1x in ice-cold PBS (0.8mM CaCl_2_) and acquired on a LSRFortessa Cell Analyzer (BD Biosciences). The experiment was performed in two independent replicates and analysed by FlowJo v10.6.1 (FlowJo LLC).

### Protein isolation, Immunoprecipitations, and Western blots

For protein extraction, hippocampi tissues were manually homogenized in 300 µl ChABC buffer (40 mM Tris-HCl pH 8.0, 40 mM sodium acetate, C3667, Sigma) containing Benzonase supplemented with Halt protease/phosphatase inhibitor cocktail (Thermo Scientific). After 10 minutes full-speed centrifugation, the supernatant containing the soluble protein fraction was separated from the pellet (insoluble fraction), which was resuspended in 300 µl ChABC buffer. Two aliquots of each fraction were brought to a protein concentration of 3 µg/µl (75µg protein in 25 µl ChABC buffer) and one aliquot of each fraction was further incubated with ChABC for 12 hours at 37°C. After adding 25 µl of Lämmli buffer, samples were heated 5 minutes at 95°C and 30µl were separated by SDS-PAGE and transferred onto PVDF membranes (Immobilion-P, Merck Millipore) according to standard protocols. Blots were blocked for 1 hour with 5% milk in TBST (1x TBS and 0.1% Tween-20) and were then incubated overnight at 4°C with primary antibodies (1:100, anti CS-0S, 1B5; anti CS-4S, 2B6, antiCs-6S, 3B3; amsbio) diluted in 5% milk in TBST. Blots were washed 3 times in TBST for 5 minutes and were then incubated with HRP-conjugated secondary anti-mouse-IgG-H&L chain (Promega) or anti-rabbit-IgG-F(ab’)2 (GE Healthcare) antibody for 1 hour at room temperature, washed 3 times in TBST for 5 minutes and visualized using enhanced chemiluminescence (ECL, GE Healthcare). See Supplementary Table 2 for a list of antibodies used in this study. β-Actin was used to control for protein loading.

#### Immunoprecipitation

N2a cells expressing FIBCD1, FIBCD1ΔFReD or empty vector were washed twice with PBS and lysed in Hunt buffer (20 mM Tris-HCl pH 8.0, 100 mM sodium chloride, 1 mM EDTA, 0.5% NP-40) supplemented with Halt protease/phosphatase inhibitor cocktail (Thermo Scientific) in 3 consecutive freeze and thaw steps. After full-speed centrifugation, the supernatant containing the soluble protein fraction was further used. An equal amount (500µg) of each protein lysate was precleared for 1 hour with magnetic Protein G Dynabeads (Invitrogen) and immunopurified with 40µl washed anti-V5 agarose beads (Sigma) overnight at 4C. After 5 washing steps in Hunt buffer, input (1/16 of the immunoprecipitation corresponding to 30µg) and immunoprecipitation samples were separated by SDS-PAGE, blotted and stained with an antibody against V5 (ab15828, 1:2000 dilution), and Western blotting was performed as described above.

### Immunofluorescence/IHC

#### Primary Neurons

At 2 or 14 days after plating (DIV2 and DIV14, respectively), primary neuronal cultures were washed with PBS and then fixed in 4% PFA with 4% glucose for 10 minutes at room temperature, and subsequently quenched with 10 µM glycine/PBS for 10 minutes at room temperature. They were then washed twice with 0.01% Triton-X/PBS (PBST), permeabilized with 0.25% Triton-X/PBS for 3 minutes and blocked in 5% NGS for 1 hour. Primary antibodies were incubated overnight at 4°C and washed 3 times in PBST. Secondary antibodies were incubated for 1 hour at room temperature and washed 3 times with PBST before mounting in ProLong Gold Antifade Mountant (Thermo #P10144). Images were acquired on fluorescent microscope (Leica), acquiring semi-random fields of the coverslips. Images were analysed manually using ImageJ. Experimenters were blinded to genotype and plating conditions during image analysis.

#### Drosophila melanogaster

Wandering third instar female larvae were dissected in Ca+ free PBS and fixed in 4% paraformaldehyde for 20 minutes. Larval body walls were blocked in serum, then incubated in primary antibodies overnight and secondary antibodies at room temperature for 2 hours while rocking. Samples were mounted in Prolong Diamond (Invitrogen) for analysis using the Zeiss LSM780 confocal microscope. Bouton number and axon branching were all identified using the anti-HRP antibody using ImageJ (NIH).

#### HEK 293T cells

Untransduced HEK 293T cells and HEK 293T cells stably expressing full length 3xFLAG tagged (C-terminal) hFIBCD1, hFIBCD1_W6*, hFIBCD1_G29S, hFIBCD1_R406C, hFIBCD1_P456L were seeded onto glass coverslips. The next day, the cells were rinsed 1x with PBS and fixed in 4% PFA (supplemented with 4% glucose) for 10 minutes at room temperature, then quenched for 10 minutes with PBS (0.01M glycine) for another 10 minutes at room temperature. The cells were washed 2x with PBS then permeabalised with 0.25% Triton-X PBS for 1.5 minutes, washed 1x with PBS and blocked for 1 hour with 5% normal goat serum at room temperature. Primary antibodies (anti-FLAG, 1:1000) were added overnight at 4°C, washed 2x with PBS, then secondary antibodies added (Alexa Fluor Anti-mouse 546, 1:500) and DAPI (1:2000) incubated at room temperature for 1 hour, washed again 2x with PBS then mounted in ProLong Gold Antifade mounting media and imaged on a Zeiss LSM980 confocal microscope.

### MRI imaging

Anaesthetised male C57Bl6/J mice (12 month of age) and their *Fibcd1* KO littermates were imaged in the Preclinical Imaging Facility at Vienna Biocentre Core Facilities (pcPHENO, VBCF), member of Vienna Biocentre (VBC), Austria, as described previously^72^. Briefly, a 15.2 Tesla MRI (Brucker BioSpec, Ettlingen Germany) was used to image brains of anaesthetised animals (isoflurane 4% induction, maintenance with 1.5%, Vana GmbH) with a 3D fast long angle shot (FLASH) sequence with magnetization transfer pulse. Each 3D image set was manually segmented using Amira 5.6 (Visualization Science Group) using The Paxinos mouse brain atlas as a reference ^73^. Values were normalized to brain size, averaged and presented as percentage of total brain volume.

### PNN visualisation with WFA

Brain was dissected from 8-10 week-old *Fibcd1* WT and KO mice and fixed in 4% paraformaldehyde and paraffin-embedded followed by 20µm coronal sectioning. For staining, the tissues were re-hydrated with PBS, permeabilized in 0.5% Triton-X PBS for 15 mins at room temperature (RT) followed by blocking in 10% goat serum diluted in 0.25% Triton-X PBS for 1 hour at RT. After 2x 3 mins washes with 0.1% tween-20 PBS, endogenous IgGs were blocked using goat F(ab) anti-mouse IgG (1:2000) diluted in 0.1% tween-20 PBS for 1 hour at RT. Next, tissues were washed 3x 3mins in 0.1% tween-20 PBS. Fluorescein *Wisteria Floribunda* Lectin (FL-1351, 1:500, Vector Laboratories) was diluted 1:500 in 0.1% tween-20 PBS with 2% goat serum and DAPI (Roth #6335, 1:2000) for 1 hour at RT. Tissues were washed 3x 3mins in 0.1% tween-20 PBS and mounted using Fluoromount Aqueous Mounting Medium (Sigma #F4680). Images were acquired using Vectra Polaris (Akoya).

### Tissue sGAG quantification

11-13-week-old *Fibcd1* WT and KO mice (mix of male and female) were sacrificed by CO_2_ asphyxiation and the hippocampi dissected out and snap frozen in liquid nitrogen. Samples were weighed and analysed for sulphated GAG (sGAG) amounts using Blyscan Glycosaminoglycan Assay kit (Biocolor #B1000), following the manufacturer’s instructions, including Papain extraction (Sigma #P3125).

### HPLC

Chondroitin sulphate was extracted from defatted, pronase digested mouse microdissected hippocampi (CA1 region) and digested using Chondroitinase ABC (from *Proteus vulgaris*, Sigma-Aldrich #C3667). The resulting glycosaminoglycan-disaccharides were fluorescently labelled with 2-aminobenzamide by reductive amination and were analysed according to previous literature^74^. Identity of glycosaminoglycan-derived disaccharides was inferred from retention time alignment with the major constituents of chondroitin sulphate sodium salt from shark cartilage (Sigma-Aldrich, C4384), and bovine trachea (Sigma-Aldrich #C9819).

### Behaviour assays

#### Mouse

Elevated Plus Maze were performed as described previously using an automated activity system (TSE-Systems, Germany)^72^. Mice were trained in the Morris Water Maze (pcPHENO, VBCF) as described previously^72^. T-Maze was performed as described previously^72^.

#### Drosophila negative geotaxis assay

Female *Nsyb-Gal4* animals were crossed with the *UAS-RNAi* lines targeting *CG10359*. Female offspring were tested at 10d after eclosion. Offspring were flipped every third day and no more than 10 flies were kept in each vial. Flies were knocked-out with CO_2_, sorted into batches of 3-7 and given 25h to recover before testing. On the day of testing, flies were flipped into empty vials and given 10-15m to recover. The climbing index is the percentage of flies that pass the 5cm mark in 5s after gently tapping to the bottom of a plastic vial.

### Acute hippocampal slice preparation and electrophysiological recordings

Memory-related synaptic plasticity and electrophysiological recordings were studied *ex vivo* in hippocampal slices essentially as previously described^75–80^. Briefly, mice were sacrificed by rapid cervical dislocation and swift sharp-blade decapitation. Brains were rapidly extracted and immediately immersed in a frosty artificial cerebrospinal fluid solution (aCSF) containing (in mM): 125 NaCl, 2.5 KCl, 25 NaHCO_3_, 2 CaCl_2_, 1 MgCl_2_, 25 D-glucose, and 1.25 NaH_2_PO_4_ (all from Sigma-Aldrich). In all experimental conditions, aCSF was continuously bubbled with a mixture of 95% oxygen and 5% carbon dioxide (Carbogen). Hippocampi were dissected while submerged in aCSF and transverse slices (300 μm in thickness) were obtained using a McIlwain Tissue Chopper (Mickle Laboratory Engineering, UK). Slices were quickly transferred to a custom-built submerged recovery chamber, where they rested for at least 1 hour submerged in 100 ml of aCSF maintained at 30 ± 2 °C. For enzymatic treatments, slices were transferred to independent home-made digestion chambers containing 15 mL of aCSF (with 0.1% bovine serum albumin) plus 0.2U/mL of either Penicillinase (Pen) (Sigma-Aldrich #61305) or Proteus vulgaris Chondroitinase-ABC (ChABC) (Sigma-Aldrich#C3667). Slices were enzymatically treated for 2 hours at 37°C. After enzymatic treatment, slices were rinsed in separated beakers containing 100 mL of aCSF (32 ± 1 °C) and gently transferred back to their corresponding recovery chamber for subsequent electrophysiological analysis. The synaptic function was studied by examining the electrical properties of the CA3-Schaffer collateral to CA1 circuit. The CA3-CA1 Schaffer collateral pathway was stimulated electrically via a home-made bipolar tungsten electrode insulated to the tip (50 µm tip diameter) and using an ISO-STIM 01D isolator stimulator (NPI Electronics, Tamm, Germany). Evoked field excitatory postsynaptic potentials (fEPSPs) were recorded at the CA1 area using aCSF-filled glass micropipettes (2– 4 MΩ) located about 400 µm away from the stimulating electrode. Input/output curves were obtained by delivering increasing pulses of voltage (100 µs in duration) between 0-9 V with a delta of 1 V and 10 s between pulses. The strength of synaptic transmission was determined in each case from the decaying slope of recorded fEPSPs. To examine paired-pulse-induced synaptic facilitation, two pulses of voltage with a strength eliciting 40% of the maximum inducible fEPSPs amplitude as determined by input/output measurements (40% fEPSP_max_) were delivered at variable inter-pulse intervals ranging between 20-100 ms with a delta increment of 20 ms (pulse pairs delivered every 10 s). The decaying slopes of the evoked fEPSPs for each consecutive pair of pulses was measured and the strength of synaptic potentiation was determined from the 2^nd^/1^st^ fEPSPs slope ratio. To study long-term potentiation, basal synaptic transmission (baseline) was examined for at least 20 min by recording stable fEPSPs in response to 40% fEPSP_max_ stimulating voltage pulses (100 µs duration; fEPSPs elicited at 0.03 Hz). After recording a steady baseline, a theta-burst stimulation (TBS) protocol was applied, consisting of five trains of 40% fEPSP_max_ stimulating voltage pulses at 100 Hz (100 µs/pulse, with 4 s inter-train interval). Postsynaptic signal in response to baseline stimulating conditions was measured for 35-70 min as indicated in figure legends. Synaptic potentiation was determined by examining the temporal course of the decaying fEPSPs slopes following TBS, normalized to baseline values. Data from fEPSPs slopes attained when measuring long term potentiation were averaged for every 2 minutes. All recordings were made using an AxoClamp-2B amplifier (Bridge mode) and a Digidata-1440 interface (Axon Instruments). Data (5-22 slices/condition) were analysed using the pClamp-10 Program software (CA/Molecular Devices, USA).

### Sequence alignment

Human and mouse Fibcd1 and Drosophila CG10359 ‘Fibrinogen C-terminal’ protein sequences were acquired from Uniprot.org. Multiple sequence alignment (MSA) was done using ClustalO algorithm (https://www.ebi.ac.uk/Tools/msa/clustalo/)81.

### Statistical analysis

#### Flow cytometry

Samples analysed by 1-way ANOVA.

#### Immunofluorescence

Images were counted manually and analysed by 1-way ANOVA. Experimenters were blinded to genotype and condition during analysis.

#### Mouse behaviour

Data was analysed by unpaired Student’s t-test. n= an individual mouse.

#### Drosophila

All data sets were organized and analysed in Microsoft Excel and Prism. Statistical test type listed in the figure legends. Data sets with equal variance were analysed using ANOVA and Dunnett’s post-hoc analysis for multiple comparison. Significance is defined as P<0.05 and error bars are shown as standard error of the mean (SEM) unless otherwise noted.

#### Electrophysiology

Two-way ANOVA with repeated measures was used to compare values between treatment groups for input/output, paired-pulse and LTP set of data as depicted in scatter charts comparing pairs of values. Two-way ANOVA was used to compare values between treatment groups at a single time or voltage point as depicted in bar charts. Data values are represented as mean ± S.E.M. (p values < 0.05 were considered statistically significant). * *p* ≤0.05; ** *p* ≤0.01; *** *p* ≤ 0.001; **** *p* ≤ 0.0001

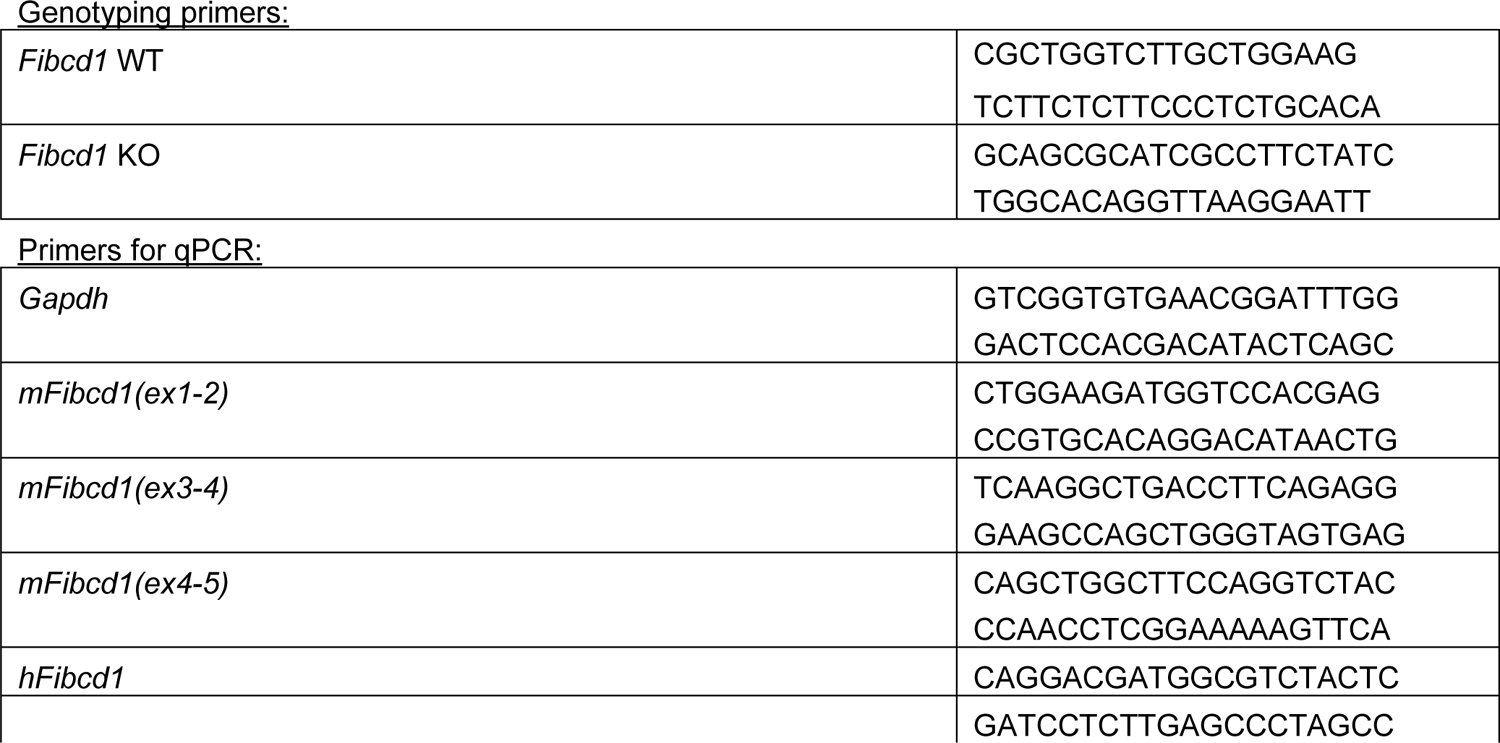
Table of oligonucleotide sequences

Table of additional materials

**Table of additional materials**
|  |  |
| --- | --- |
| <u>Antibodies for immunoblots:</u> |  |
| β-Actin | A5316 (Sigma) |
| CS-0S (1B5) | 270431-CS (Amsbio) |
| CS-4S (2B6) | 270432-CS (Amsbio) |
| CS-6S (3B3) | 270433-CS (Amsbio) |
| Anti-V5 tag | Ab15828 (Abcam) |
| <u>Fluorescent Sugars for Flow Cytometry:</u> |  |
| Fluoresceinamine Labeled Sodium Chondroitin Sulfate A (A1) | AMS.CSR-FACS-A1 (Amsbio) |
| Fluoresceinamine Labeled Sodium Chondroitin Poly-Sulfate (P1) | AMS.CSR-FACS-P1 (Amsbio) |
| Fluoresceinamine Labeled Sodium Dermatan Sulfate (B1) | AMS.CSR-FADS-B1 (Amsbio) |
| <u>Antibodies/dyes for immunofluorescence:</u> |  |
| MAP2 | Millipore 05-346 |
| FLAG (M2) | Sigma F1804 |
| Alexa Fluor® 546 anti-mouse | Thermo A-11003 |
| Goat F(ab) anti mouse (IgG) | Abcam (ab6668) |
| Alexa Fluor® 647 AffiniPure Goat Anti-Horseradish Peroxidase | Jackson Immunoresearch |
| Mouse anti-nc82 (Bruchpilot) | Developmental Hybridoma Studies Bank |
| DAPI | Carl Roth |
| WFA-488 | Vector Laboratories (FL-1351) |

Table of Drosophila Reagents

**Table of Drosophila Reagents**
|  |
| --- |
| <u>Drosophila reagents</u> |

| <u>Stock</u> | <u>RRID/source</u> |
| --- | --- |
| y[1] w[*]; P{w [+mC]=r4-GAL4}3 | BDSC_33832 |
| y[1] v[1]; P{y[+t7.7] v[+t1.8]=TRiP.HMJ30271}attP40 | BDSC_63703 |
| w1118; P{GD2280}v4128/TM3 | FlyBase_FBst0464025 |
| P{KK105143}VIE-260B | FlyBase_FBst0474536 |
| y[1] w[*]; P{w[+m*]=nSyb-GAL4.S}3 | BDSC_51635 |
| y[1] w[*]; P{w[+mC]=tubP-GAL4}LL7/TM3, Sb[1] Ser[1] | BDSC_5138 |
| PBac{UAS-empty}VK00037 | Yoon et al., Cell Metabolism, 2017 |
| y[1] v[1]; P{TRiP.JF01355}attP2 | BDSC_31603 |

## SUPPLEMENTARY FIGURE TITLES AND LEGENDS

**Figure S1:**
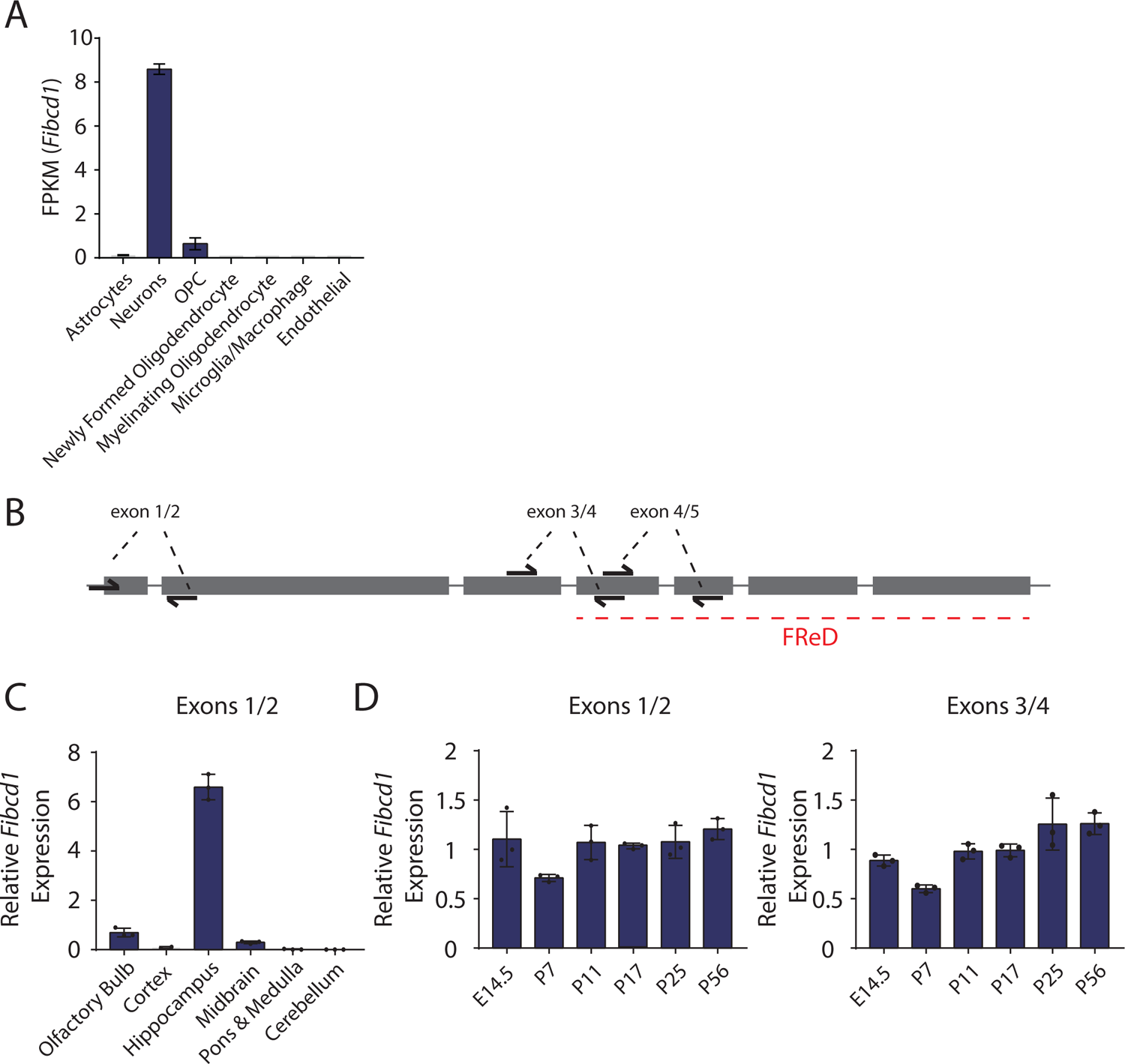
*Fibcd1* expression in the adult and developing mouse brain. (A) *Fibcd1* expression in bulk populations of sorted mouse brain cell population, from *brainrnaseq.org.* OPC = oligodendrocyte precursor cell. (B) Schematic of *Fibcd1* exons (grey rectangles) and introns (grey lines), and location of primer pair binding (‘exons 1/2, 3/4 and 4/5’) used for RT-qPCR. Exon sizes are to scale; introns and primer lengths are not. The exons coding for FIBCD1 FReD is indicated by a red dashed line. (C) Relative mRNA expression levels of *Fibcd1* (primers binding to exon 1 and 2) normalised to *Gapdh* in the indicated brain regions, analysed by RT-qPCR (N = 3). (D) Relative mRNA expression levels of *Fibcd1* (primers binding to exon 1 and 2 and exons 3 and 4) normalised to *Gapdh* in the hippocampus of the indicated time points, analysed by RT-qPCR (N = 3). Error bars represent SD.

**Figure S2:**
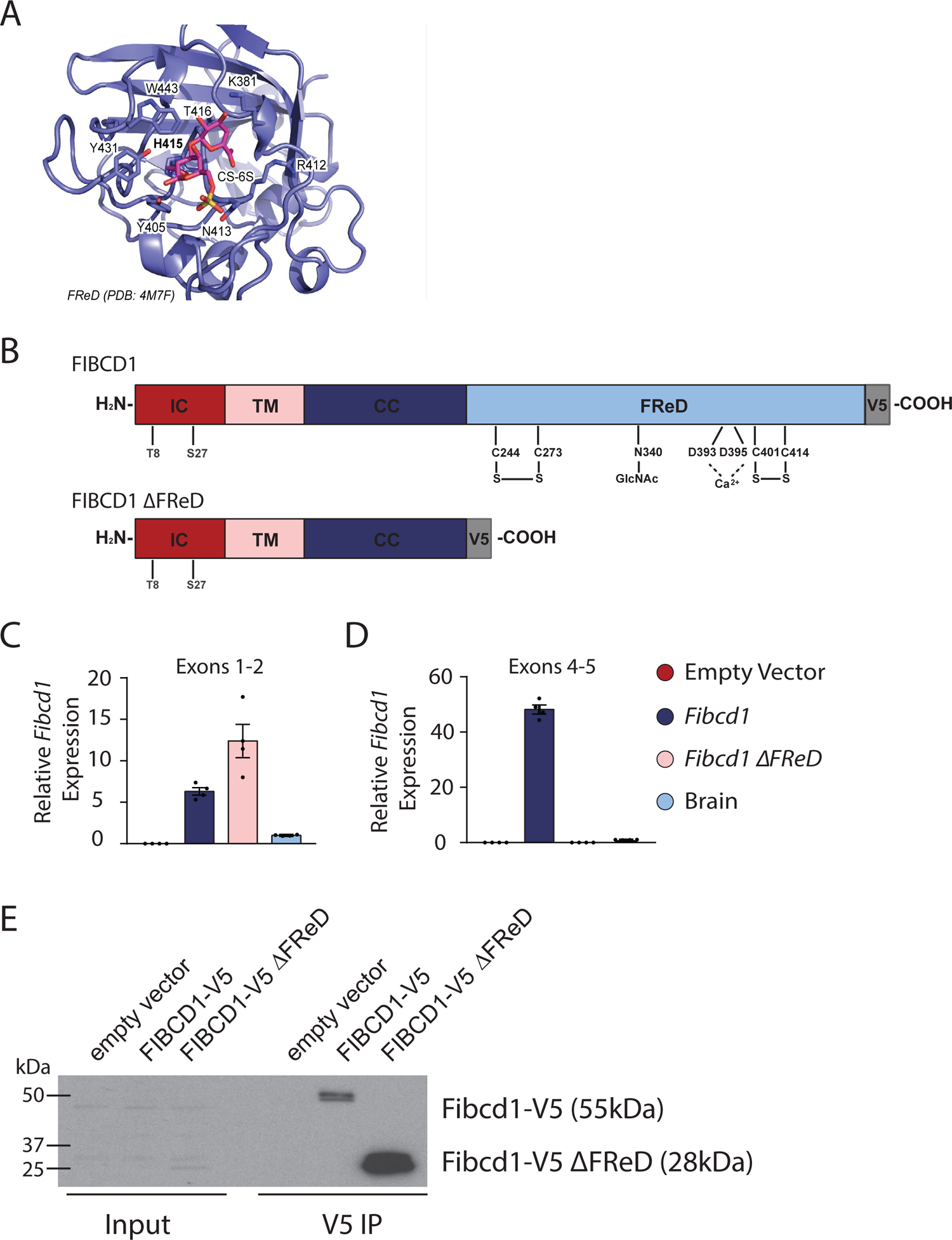
Docking site of CS-6S in FIBCD1 FReD and validation of mFIBCD1 overexpressing N2a cell lines. (A) Top binding pose for *in silico* docking of CS-6S to FIBCD1 FReD (PDB 4M7F). (B) Schematic representation of FIBCD1 domains, IC-intracellular domain (red), TM-transmembrane domain (pink), CC-coiled coil domain (dark blue), FReD (light blue), and location of V5-tag (grey) in full-length mFIBCD1 cDNA and truncated mFIBCD1 lacking the FReD domain (FIBCD1 ΔFReD). (C) Relative mRNA expression levels of *Fibcd1* in the N2a cells overexpressing full-length (*Fibcd1*) or truncated *FIBCD1* (*Fibcd1 ΔFReD*) and adult mouse WT brain for comparison, analysed by RT-qPCR (n = 2). Primers binding to exon 1 and 2 before the FReD domain (B) or to exon 4 and 5 spanning the sequence encoding part of the FReD. Note the complete absence of endogenous *Fibcd1* expression in the ‘empty vector’ (red bar) control and the complete absence of expression when using primers against exon 4/5 (D), which span the FReD (see Figure S1B) in the *Fibcd1* ΔFReD construct (C, pink bar), validating the generated cell lines. *Gapdh* was used as housekeeping control and values obtained from a control brain sample were set to 1. (E) Validation of transgenic N2a cell line at the protein level by immunoprecipitation with anti-V5 antibody as bait. Input (left) and V5-immunoprecipitated (right) lysates from N2a cells expressing V5-tagged full-length mFIBCD1 (mFIBCD1-V5, predicted size of 55kDa), V5-tagged mFIBCD1 lacking the FReD (V5-FIBCD1 ΔFReD, predicted size of 28kDa) or the empty vector as negative control. Protein marker sizes are indicated.

**Figure S3:**
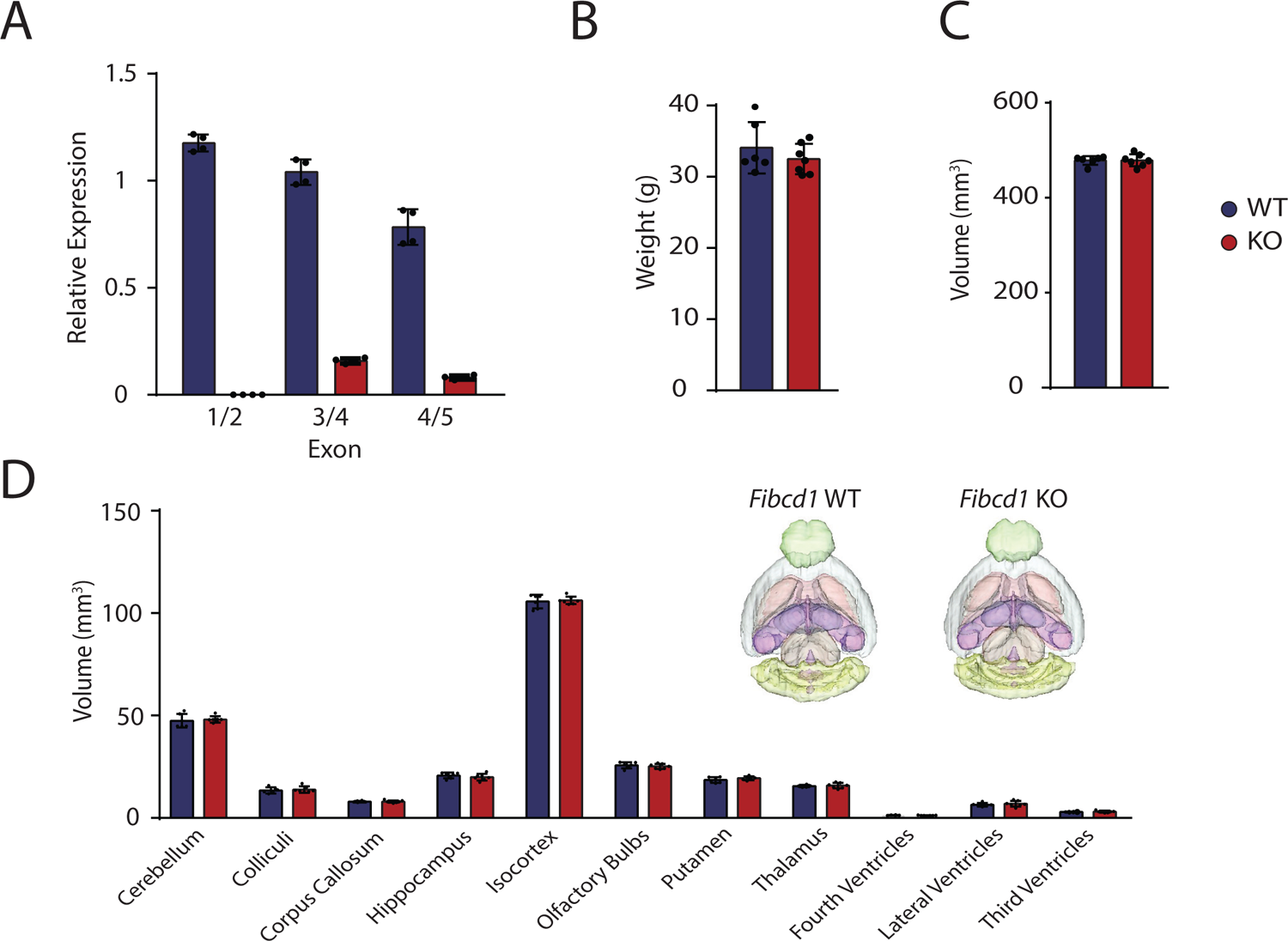
*Fibcd1* expression, weight and brain tissue volumes in *Fibcd1* WT and KO mice. (A) RT-qPCR of *Fibcd1* WT and KO adult mouse hippocampi using primer pairs binding to indicated exons (see Fig. S1B). N = 4. (B-D) Body weight (B), total brain volume (C) and brain volumes of denoted brain regions (D) of the indicated genotypes as assessed by MRI volumetric analysis (N > 5). Inset are 3D representative MRI renditions of control (left) and *Fibcd1* KO (right) adult brains with analysed brain regions pseudo-coloured. Error bars represent SD.

**Figure S4:**
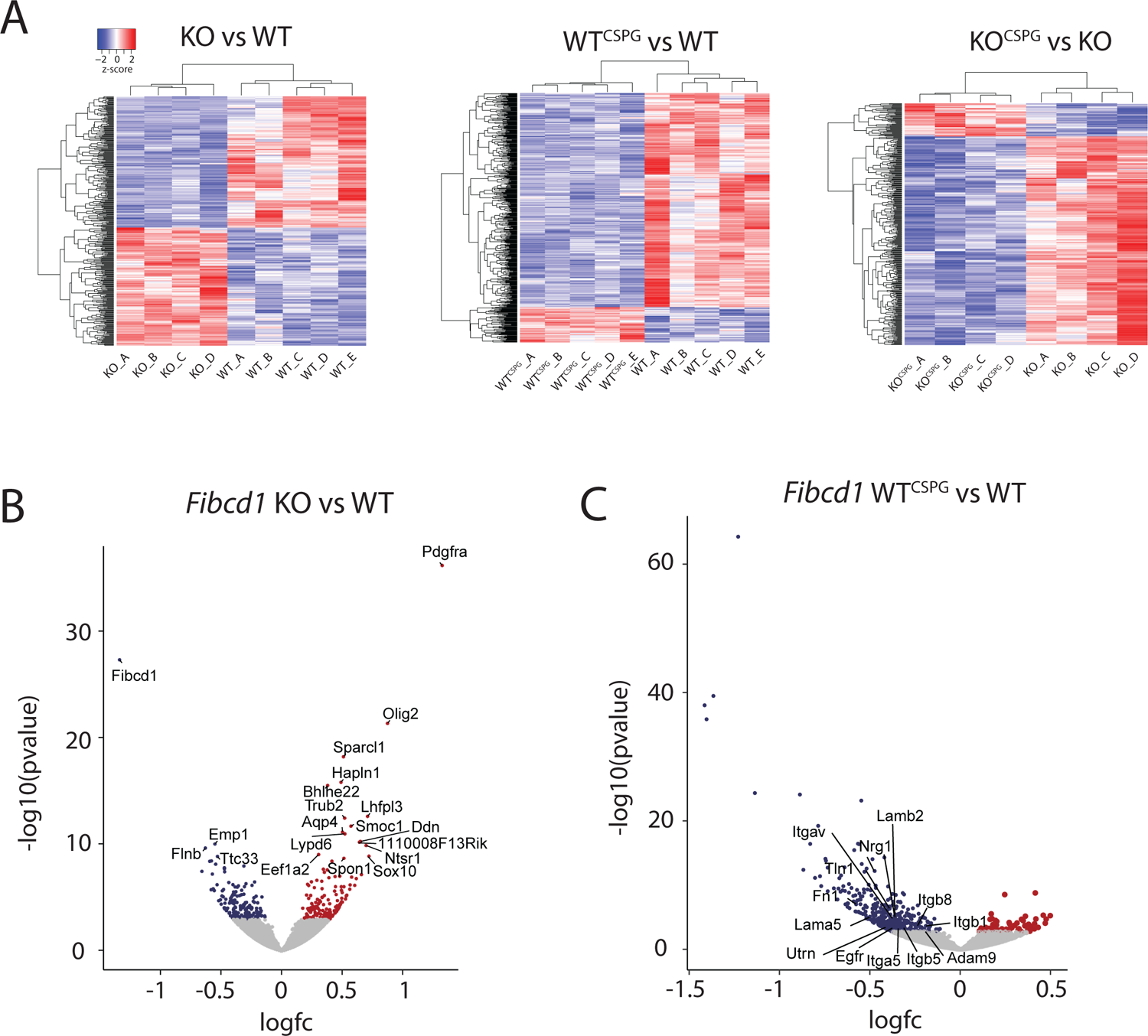
Additional RNA-seq analysis. (A) z-score hierarchical clustering for each sample in *Fibcd1* KO vs WT, WT^CSPG^ vs WT and KO^CSPG^ vs KO. Colours represent scaled expression values, with blue for low and red for high expression levels. Legend is indicated. (B) Volcano plot depicting differential gene expression at DIV2 hippocampal cultures, comparing *Fibcd1* KO vs WT, showing significantly upregulated (red) and downregulated (blue) genes. Top 20 differentially expressed genes are labelled. (C) Volcano plot depicting differential gene expression at DIV2 hippocampal cultures comparing WT^CSPG^ vs WT, all genes that fall into “integrin binding” GO term category are labelled.

**Figure S5:**
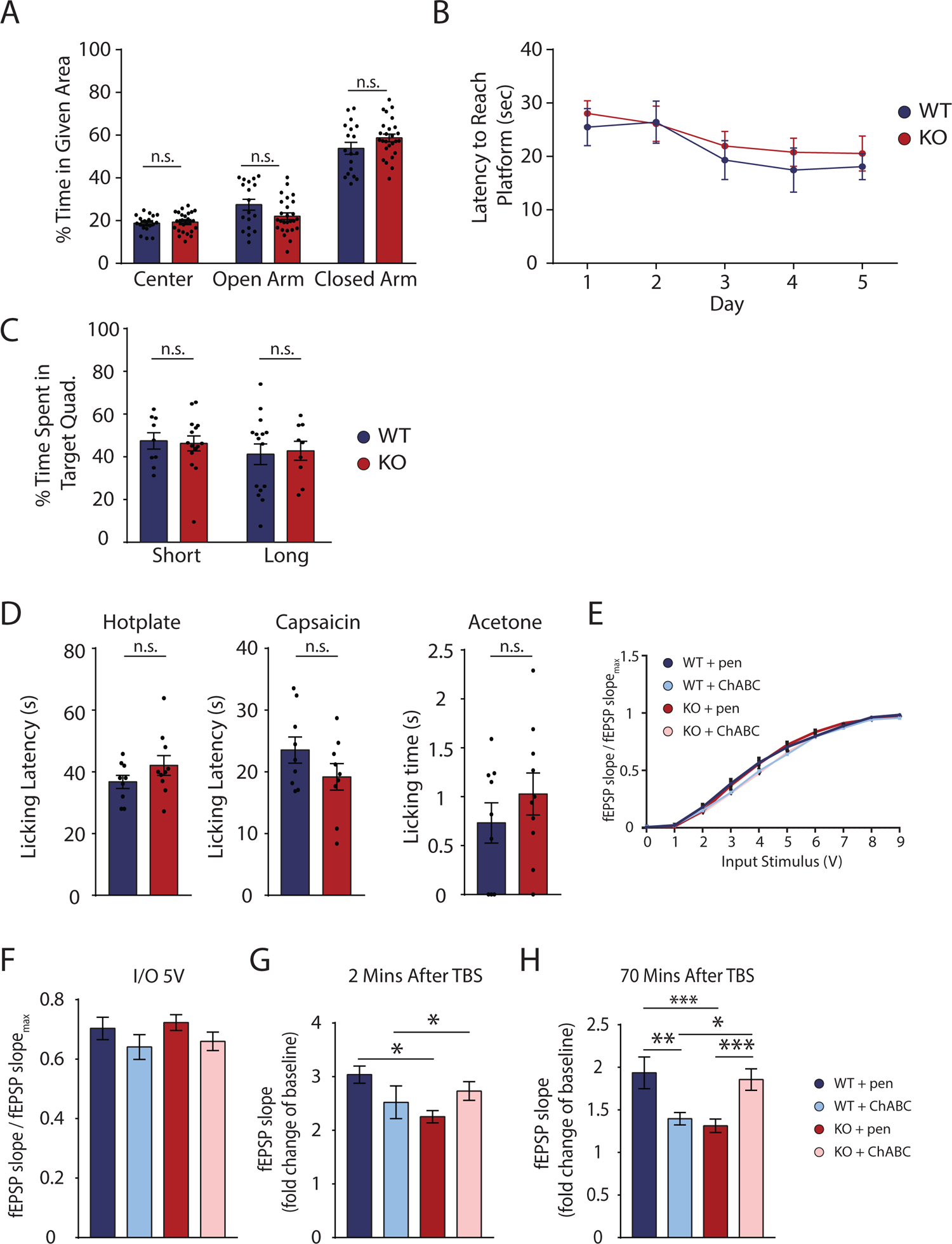
Analysis of *Fibcd1* KO mouse neurological phenotype. (A) Percentage of time mice spent in the centre, open and closed arms of the Elevated Plus Maze (EPM) are shown for *Fibcd1* WT (blue) and KO (red). N = 18 WT; 27 KO. (B-C) Mouse performance quantified by time to reach the target platform (B) and time spent in the target quadrant (C), in Morris Water Maze (MWM) hippocampal-dependent spatial learning task over 5 days. N = 5 WT; 9 KO. (D) Acute pain responses to hotplate, acetone drop, or intraplantar capsaicin injections, quantified as time to first response or time spent licking or biting the injected paw, respectively. N= 9 WT; N = 10KO. (E-F) Input/output assessment of synaptic transmission in CA3-CA1 Schaffer collateral pathway of adult mouse hippocampal slices. *Fibcd1* WT (blue) and KO (red) hippocampal slices, pre-treated with penicillinase (pen) or Chondroitinase ABC (ChABC). N (WT+pen) = 22; N (KO+pen) = 27; N (WT+ChABC) = 21; N (KO+ChABC) = 30. (G-H) LTP fold change of baseline at 2 (G) and 70 (H) minutes post theta-burst stimulation (TBS) in adult mouse hippocampal slices. N (WT+pen) = 9; N (KO+pen) = 15; N (WT+ChABC) = 6; N (KO+ChABC) = 12. Not significant, n.s., * = p≤0.05; ** =p≤0.01; *** =p≤0.001. Error bars represent SEM.

**Figure S6:**
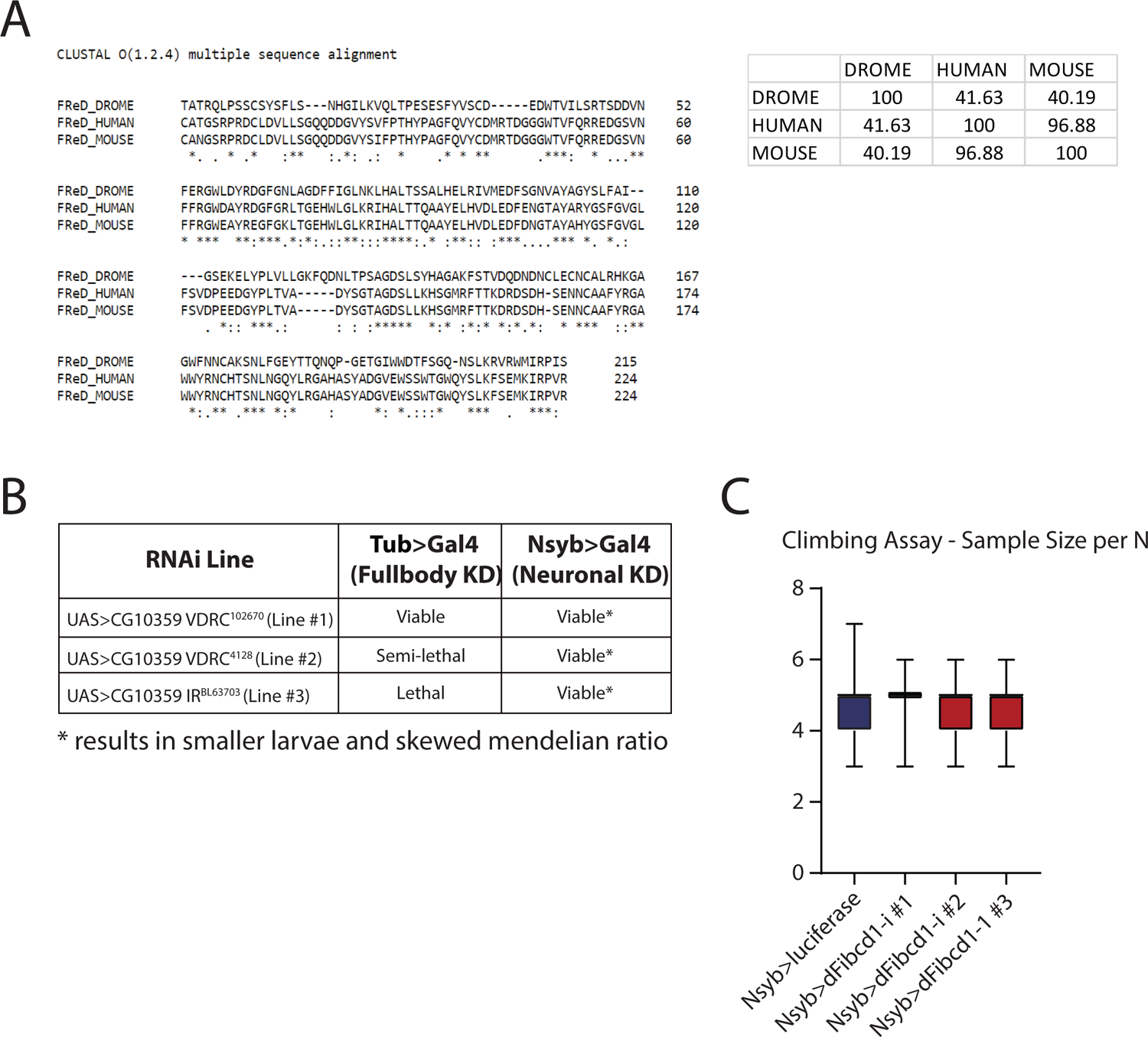
Description of *dFibcd1*. (A) Alignment of human, mouse and fly (DROME) FReD protein sequence. Inset, percent identity matrix (% homology) between Drosophila, human and mouse FReD protein sequences. (B) Summary of 3 RNAi lines crossed to full body GAL4 driver (tubulin) or neuron-specific (Nsyb) and the effects on viability. (C) Number of flies analysed for the climbing assay in Fig 4I.

**Figure S7:**
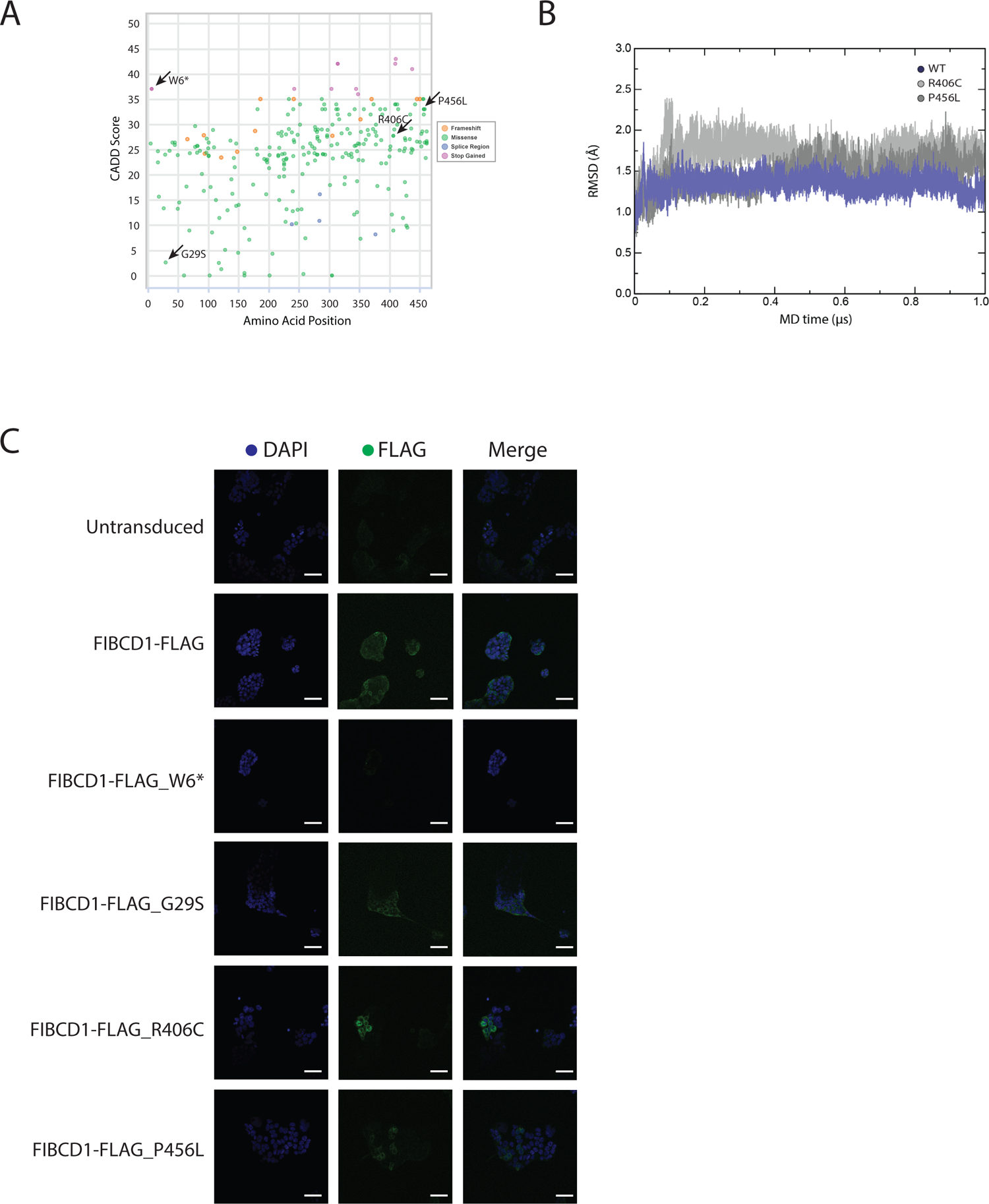
Additional human Fibcd1 data. (A) Missense, frameshift, splice region and stop gain variants present in the population. Each dot represents one distinct variants, amino acid position and CADD score indicated on x and y axis. Indicated with arrows are the variants discussed in the present study. Data originally from gnomAD. (B) Time course of the backbone RMSD from the starting configuration for WT (blue), R406C (pink) and P456L (red) MD simulations. (C) Validation of FIBCD1 expression in stably expressing HEK293t cells by immunofluorescence. Shown is DAPI (blue), anti-FLAG (green) and merge. Scale bar = 50um.

